# Getting close to nature – *Plasmodium knowlesi* reference genome sequences from contemporary clinical isolates

**DOI:** 10.1101/2021.11.16.468780

**Authors:** Damilola R. Oresegun, Peter Thorpe, Ernest Diez Benavente, Susana Campino, Fauzi Muh, Robert Moon, Taane G. Clark, Janet Cox-Singh

## Abstract

*Plasmodium knowlesi,* a malaria parasite of old-world macaque monkeys, is used extensively to model *Plasmodium* biology. Recently *P. knowlesi* was found in the human population of Southeast Asia, particularly Malaysia. *P. knowlesi* causes un-complicated to severe and fatal malaria in the human host with features in common with the more prevalent and virulent malaria caused by *Plasmodium falciparum*.

As such *P. knowlesi* presents a unique opportunity to inform an experimental model for malaria with clinical data from same-species human infections.

Experimental lines of *P. knowlesi* represent well characterised genetically static parasites and to maximise their utility as a backdrop for understanding malaria pathophysiology, genetically diverse contemporary clinical isolates, essentially wild-type, require comparable characterization.

The Oxford Nanopore PCR-free long-read sequencing platform was used to sequence *P. knowlesi* parasites from archived clinical samples. The sequencing platform and assembly pipeline was designed to facilitate capturing data on important multiple gene families, including the *P. knowlesi schizont-infected cell agglutination* (*SICA*) *var* genes and the *Knowlesi-Interspersed Repeats* (*KIR*) genes.

The *SICAvar* and *KIR* gene families code for antigenically variant proteins that have been difficult to resolve and characterise. Analyses presented here suggest that the family members have arisen through a process of gene duplication, selection pressure and variation. Highly evolving genes tend to be located proximal to genetic elements that drive change rather than regions that support core gene conservation. For example, the virulence-associated *P. falciparum* erythrocyte membrane protein (*PfEMP1*) gene family members are restricted to relatively unstable sub-telomeric regions. In contrast the *SICAvar* and *KIR* genes are located throughout the genome but as the study presented here shows, they occupy otherwise gene-sparse chromosomal locations.

The novel methods presented here offer the malaria research community new tools to generate comprehensive genome sequence data from small clinical samples and renewed insight into these complex real-world parasites.

**Author summary:** Malaria is a potentially severe disease caused by parasite species within genus Plasmodium. Even though the number of cases is in decline there were over 200 million reported cases of malaria in 2019 that resulted in >400,000 deaths. Despite huge research efforts we still do not understand precisely how malaria makes some individuals very ill and by extension how to successfully augment and manage severe disease.

Here we developed a novel method to generate comprehensive robust genome sequences from the malaria parasite *Plasmodium knowlesi* collected from clinical samples.

We propose to use the method and initial data generated here to begin to build a resource to identify disease associated genetic traits of *P. knowlesi* taken from patient’s samples. In addition to the methodology, what further sets this work apart is the unique opportunity to utilize same-species experimental *P. knowlesi* parasites to discover a potential role for particular parasite traits in the differential disease progression we observe in patients with *P. knowlesi* malaria.

While we developed the methods to study severe malaria, they are affordable and accessible, and offer the wider malaria research community the means to add context and insight into real-world malaria parasites.

## Introduction

*Plasmodium knowlesi* is a malaria parasite first described in a natural host, the long-tailed macaque monkey (*Macaca fascicularis*), in the early part of the 20^th^ Century [1]. Although an incidental find, *P. knowlesi* was soon exploited as a model parasite for malaria research [2–4]. Experimental *P. knowlesi* was well-characterised over time with several additional lines adapted from natural hosts in geographically distinct regions, including a human infection [2–6]. Taken together, experimental lines of *P. knowlesi* are important members of the malaria research arsenal.

What sets *P. knowlesi* apart is that it occupies several important niche areas - as an experimental model, a natural parasite of Southeast Asian macaque monkeys and the causative agent of zoonotic malaria in the human host [7]. In nature, transmission is established in the jungles of Southeast Asia, areas that support the sylvan mosquito vectors, the parasite and the natural macaque hosts. People who enter transmission sites are susceptible to infected mosquito bites and infection. *P. knowlesi* has effectively crossed the vertebrate host species divide and is responsible for malaria in contemporary human hosts [8].

Zoonotic malaria caused by *P. knowlesi* is currently the most common type of malaria in Malaysia, with most of the cases reported in Malaysian Borneo [9]. Indeed, naturally acquired *P. knowlesi* malaria causes a spectrum of disease from uncomplicated to severe and fatal infections with tantalizing similarity to severe adult malaria caused by *P. falciparum* [10–13]

The clinical similarities observed in patients with severe *P. knowlesi* and *P. falciparum* infections suggest that *P. knowlesi* has the potential to serve as both a human pathogen and animal model for severe malaria pathophysiology that has hitherto eluded medical science [11, 14, 15].

To take this idea forward, it seemed prudent to compare genome sequences derived from contemporary clinical isolates of *P. knowlesi* with the reference *P. knowlesi* genome generated from a genetically static and laboratory passaged experimental line [16].

We developed methods to produce high-quality Illumina short-read *P. knowlesi* genome sequence data from frozen clinical blood samples [17]. The outputs of this work identified genome-wide diversity, including genomic dimorphism in *P. knowlesi* isolates from patients. Comparisons also highlighted that reference *P. knowlesi* genome sequence data, generated from experimental lines established mid-twentieth century, may not properly reflect and capture important loci for research on malaria pathophysiology, particularly multiple gene families.

*Plasmodium* species have a number of multiple gene families encoding infected red blood cell surface proteins that are antigenic and highly variable to avoid host immune recognition and parasite destruction [18, 19]. Of these are the *P. falciparum* erythrocyte membrane protein (*PfEMP1)* gene family members with an estimated 67 copies in the *P. falciparum* 3D7 reference genome and variable copy numbers in clinical isolates (n = 47 – 90) [20] [21]. While other multiple gene families are described in all *Plasmodium* species studied to date, *PfEMP1* gene-like families are rare, and among the parasites that cause human infection, are found only in *P. falciparum* and *P. knowlesi* [16, 20]. *PfEMP1* genes are expressed in a mutually exclusive manner with only one predominantly expressed at any one time [22–24]. Importantly *PfEMP1* gene expression is implicated in *P. falciparum* virulence and progression to severe disease [19, 22, 25–29]. The comparable *P. knowlesi schizont-infected cell agglutination* (*SICA*) *var* gene family has been reported in detail in various experimental lines [3, 16, 30, 31]. Corredor *et al*, (2004) described conserved yet polymorphic repeat patterns in a 3’ untranslated region (*SICAvar* 3’ UTR sequences) of a particular SICA gene from the experimental clone *P. knowlesi* Pk1B^+^. They suggest the *SICAvar* 3’ UTR may be a site for extensive recombination and have implication in post-transcriptional *SICAvar* gene expression regulation [30–32]. To our knowledge, the *P. knowlesi* SICAvar gene family and 3’ UTR’s have not yet been described in-depth, in wild-type isolates, including *P. knowlesi* isolated from clinical infections. Given the *PfEMP1* gene association with severe disease in *P. falciparum,* we are particularly interested in characterising variation and disease association between the *P. knowlesi SICAvar* gene family members in clinical isolates.

Genome sequence data for multiple gene families in general and *Plasmodium* spp. in particular are difficult to resolve using Illumina short-read sequencing platforms. This is due to sequence similarity between the family members and long stretches of regions of low complexity [17] [33]. In addition, most *Plasmodium* reference genome sequences are derived from experimental lines that may incompletely represent multiple gene families. Recently, the PacBio long-read sequencing platform was used to describe, for the first time, the core *P. falciparum* genome in clinical isolates and demark sub-telomeric regions to compare genome organisation and diversity between clinical isolates from different geographical regions and the commonly used *P. falciparum* clone 3D7 [21].

Keeping in mind comparative biology, pathobiology and genomics, we propose to describe multiple gene family organisation, location and copy number in *P. knowlesi* clinical isolates using long read amplification-free sequencing. The PacBio platform is outside of our reach because we have small volume frozen whole blood samples that yield parasite DNA well below the quantity required for amplification-free PacBio sequencing [21, 30, 34]. Here we use the accessible, portable and affordable Oxford Nanopore Technologies MinION long-read sequencing platform to *de novo* assemble two new *P. knowlesi* reference genome sequences representing each of genetically dimorphic forms of *P. knowlesi* found in our patient cohort [17] [35].

The new reference genomes will, for the first time, provide insight into clinically relevant contemporary *P. knowlesi* parasites. These diverse parasites are essentially wild-type and the product of ongoing mosquito transmission and recombination in nature [17, 36–39]. The genomes will offer a valuable resource not only for our studies on members of the *SICAvar* gene family and virulence but also to the wider zoonotic malaria research community working on comparative biology of malaria parasites, drug discovery and vaccine development.

## Results

### Evaluating draft de novo genomes

The genome pipeline, beginning with Oxford Nanopore Technologies (ONT) MinION sequencing through to *de novo* assembly and genome annotation with downstream analyses, is shown (Fig 1). The pipeline was used to produce draft *P. knowlesi de novo* genomes using DNA extracted from two clinical isolates sks047 and sks048 and for comparison the well characterised cultured line, *P. knowlesi* A1-H.1. For purpose of clarity, the *P. knowlesi* A1-H.1 *de novo* draft genome assembled here is referred to as StAPkA1H1 (please see methods section). Read coverage of 225x, 71x and 65x was obtained for StAPkA1H1, sks047 and sks048 respectively (**Error! Reference source not found.**). The draft assemblies resolved into 100 or fewer contigs before further reduction to <72 contigs after scaffolding (Table 1). The quality of the draft assemblies was improved with Medaka’s polishing resulting in Benchmarking Universal Single-Copy Orthologues (BUSCO) scores that increased from 68.6 to 89.7 (a 30.8% increase), 67.2 to 85.5 (a 27.2% increase) and 68.8 to 85.9 (a 24.8% increase) for StAPkA1H1, sks047 and sks048 respectively with BUSCO completeness scores for the clinical isolates reaching 95%. (Table 1). The observed increase in the number of contigs from 23.57 to 23.63Mb (0.22% increase) for sks047 and 24.49 to 24.56Mb (0.32% increase) for sks048 was likely due to the addition of relatively shorter reads (Table 1).

**Fig 1.**
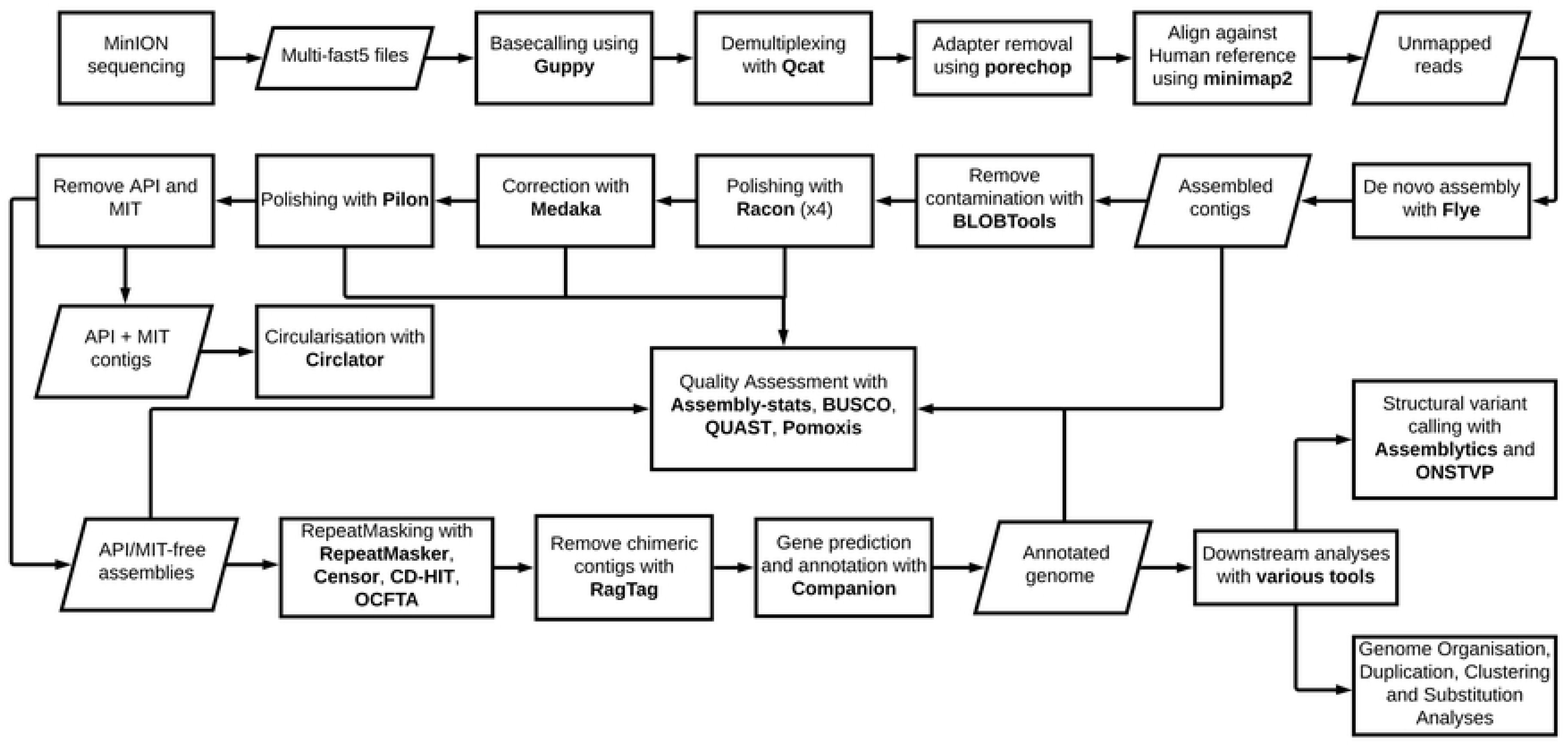
*Plasmodium knowlesi de novo* genome pipeline. The pipeline represents major forms of manipulation taken and tools utilised to generate, annotate and analyse the two reference genomes derived from clinical isolates.

**Table 1.** Overview of assembly and quality metrics of the de novo assembled draft assemblies.

| *Isolate* | *Coverage* | *de novo assembly length (Mb)* |  |  |  |  | *Contigs/Scaffolds/Chromosomes* |  |  |  |  | *BUSCO Completeness Score (%)* |  |  |  |  |
| --- | --- | --- | --- | --- | --- | --- | --- | --- | --- | --- | --- | --- | --- | --- | --- | --- |
|  |  | *Raw* | *Medaka* | *Pilon* | *RagTag* | *Complete* | *Raw* | *Medaka* | *Pilon* | *RagTag* | *Complete* | *Raw* | *Medaka* | *Pilon* | *RagTag* | *Complete* |
| *PKNH [16]* | - | - | - | - | - | 24.36 | - | - | - | - | 15 | - | - | - | - | 97.6 |
| *PKA1H1*<br>*[40]* | - | - | - | - | - | 24.27 | - | - | - | - | 14 | - | - | - | - | 94.4 |
| *StAPkA1H1* | 225X | 24.15 | 24.14 | N/A | 24.39 | 24.39 | 73 | 111 | N/A | 71 | 15 | 68.6 | 89.7 | - | 89.7 | 89.5 |
| *sks047* | 71X | 23.57 | 23.63 | 23.64 | 24.17 | 24.17 | 100 | 116 | 116 | 69 | 15 | 67.2 | 85.5 | 95.7 | 95.9 | 95.9 |
| *sks048* | 65X | 24.49 | 24.56 | 24.57 | 24.81 | 24.81 | 74 | 94 | 94 | 50 | 15 | 68.8 | 85.9 | 95.7 | 95.7 | 95.6 |

The combination of previously sequenced Illumina reads data with 34x and 166x short read coverage for sks047 and sks048 respectively offered the opportunity for Pilon polishing the newly generated ONT sequence data for clinical isolates sks047 and sks048. Pilon polishing resulted in improved BUSCO scores with sks047 seeing an 11.9% improvement (85.5 to 95.7) and sks048 showing an 11.4% improvement (85.9 to 95.7) (Table 1). Although Pilon did not change the number of contigs both sks047 and sks048 saw a total length increase of 0.05% and BUSCO score increases. Additional Illumina sequencing was not available for StAPkA1H1 and Pilon polishing was not possible.

Scaffolding, chromosome structuring and subsequent annotation initially proved difficult due to large sections of chromosomes 2 and 3 consistently being incorrectly placed in chromosomes 14 and 13, respectively. These large-scale inconsistencies were the result of contig chimers and were minimised or entirely corrected by de-chimerisation using RagTag. Chromosomes corrected by RagTag retained regions of variability for the nuclear genome assemblies (apicomplast (API) and mitochondria (MIT)-free) although RagTag did not provide a complete solution in resolving all variable sequences (SI Fig 1; SI Fig 2). In addition, it is possible that RagTag did not entirely retain highly variable regions such as telomeric regions that may have resulted in loss of coverage of genes positioned at extreme chromosomal boundaries (SI Fig 1).

### Genome Annotation and gene content

Companion software resolved all three nuclear genomes StAPkA1H1, sks047 and sks048 into 15 chromosomes – 14 Pk chromosomes and 1 ‘bin’ ‘00’ chromosome holding sequence fragments which could not be confidently placed by the Companion pipeline (Table 2). Each draft genome was assigned a similar or greater number of coding genes than the PKNH reference genome (5327 genes) when full protein-coding genes and pseudogenes annotated with predicted function (implying missing ‘start’ and/or ‘stop’ codons) were combined. The StAPkA1H1 draft assembly was found to have 5358 genes (4385 coding + 973 pseudogenes), while the patient isolate draft genomes - sks047 and sks048 had 5327 genes (4886 coding + 441 pseudogenes) and 5398 genes (4904 coding + 494 pseudogenes) respectively (**Error! Reference source not found.**). Non-coding genes were also found in all three draft genomes, including multiple small nuclear RNA (snRNA) (Supplementary File 1).

**Table 2.** Summary of the complete de novo draft genomes compared to the published *P. knowlesi* PKNH and PkA1H1 reference genomes.

| *Isolate* | *Complete*<br>*assembly*<br>*length*<br>*(Mb)\** | *Contigs* | *Chromo*<br>*somes* | *N50*<br>*(Mb)* | *N count* | *Gaps* | *Genes*<br>*\*\** | *Total*<br>*pseudo-*<br>*genes* | *Shared*<br>*Orthologous*<br>*clusters w.*<br>*reference* | *Unique*<br>*orthologous*<br>*clusters* | *Singleton*<br>*clusters* | *KIRs* | *SICAvars\*\*\** |  |  |
| --- | --- | --- | --- | --- | --- | --- | --- | --- | --- | --- | --- | --- | --- | --- | --- |
|  |  |  |  |  |  |  |  |  |  |  |  |  | T1 | T2 | SDM's |
| *PKNH [16]* | 24.36 | - | 15 | 2.16 | 11381 | 98 | 5327 | 12 | - | - | - | 61 | 89 | 20 | 127 |
| *PKA1H1[40]* | 24.27 | 156 | 14 | 2.19 | 148255 | 142 | - | - | - | - | - | - | - | - | - |
| *StAPkA1H1* | 24.39 | 71 | 15 | 2.13 | 288598 | 127 | 5358 | 973 | 4172 | 3 | 62 | 51 | 191 | 15 | 88 |
| *sks047* | 24.17 | 69 | 15 | 2.09 | 544896 | 109 | 5327 | 441 | 4666 | 9 | 82 | 22 | 115 | 9 | 181 |
| *sks048* | 24.81 | 50 | 15 | 2.21 | 283076 | 84 | 5398 | 494 | 4664 | 11 | 100 | 25 | 153 | 7 | 196 |

\* total genome length excluding the mitochondrial and apicoplast genome sequences
\*\* total number of coding genes and pseudogenes identified with a function
\*\*\* *SICAvAr* Type 1 (T1); *SICAvAr* Type 2 (T2); *SICAvAr* single domain fragments (SDM's). Single domain fragments code for *SICAvAr* protein fragments.

*Schizont-infected cell agglutination* (*SICAvar*) and the *Knowlesi-Interspersed Repeats* (*KIR*) multiple gene families were annotated in each draft genome (Table 2). There were consistently fewer *KIR* gene family members in the draft genomes derived from clinical isolates; sks047, *KIR* n = 22 and sks048 *KIR* n = 25 compared with the experimental lines StAPkA1H1 *KIR* n = 51, and the published PKNH reference genome *KIR* n = 61 (**Error! Reference source not found.**). It is unlikely that this is a result of assembly error given that StAPkA1H1 and the clinical isolates sks047 and sks048 were sequenced and *de novo* assembled in parallel using the same methodologies with the exception of Pilon polishing for StAPkA1H1. Indeed, the dN/dS ratio (see below) supports divergence of the *KIR* gene family.

All three draft genomes had more *SICAvar* Type 1 genes annotated (StAPkA1H1, *SICAvar* type 1 n = 191; sks047 *SICAvar* type 1, n =115 and sks048 *SICAvar* type 1 n = 153 compared with the reference genome PKNH *SICAvar* type 1 n = 89 (Table 2*). SICAvar* gene fragments in each of the clinical isolate draft genomes, sks047 and sks048, outnumbered annotated Type 1 genes (Table 2). Conversely the StAPkA1H1 draft genome had approximately half the number of *SICAvar* gene fragments compared with the clinical isolates and compared with StAPkA1H1 Type 1 genes (Table 2). The complement of *SICAvar* genes and gene fragments in the draft genomes presented here were resolved to the best current sequencing technology. The differences observed between *SICAvar* gene copy numbers and fragment copy numbers in clinical isolates compared with those in experimental lines deserves further investigation.

In regions of the draft genomes where gaps could not be resolved contigs which had evidence that they belong together either by long reads spanning them, or similarity to the reference, were scaffolded with N bases, proportional to the gap size (Table 2). Higher N counts are observed in the three AM-F draft genomes generated here compared with the published reference genome (PKNH). Sequences placed in the draft genome ‘bin’ chr 00 may reflect the higher N counts in chromosomes 1 – 14. The ‘bin’ chr 00 of StAPkA1H1 clustered with the PKNH reference ‘bin’ chr 00 (SI Fig 2A) suggesting the StAPkA1H1 draft genome had a similar structure to the PKNH reference genome, including ‘unplaced’ genes. In contrast, sks047 and sks048 ‘00’ chromosome sequences are distributed across the reference genomes, suggesting no single chromosome was more challenging to scaffold after de-chimerisation (SI Fig 2ii, iii). The number of gaps in the three draft AM-F draft genomes was variable but within the range of the PKNH reference genome (Table 2).

Orthologous genes were determined using a similarity approach by OrthoMCL in Companion show all three AM-F draft genomes share >4000 orthologs with the PKNH reference genome (Table 2). These orthologous genes can be considered as the core *P. knowlesi* gene set and are indicative of reliable and accurate assemblies (Table 2). In particular, the contemporary patient isolates – sks047 and sks048 – show >4600 shared orthologues with the PKNH reference genome (Table 2).

### Apicoplast and Mitochondrial circularisation

The apicoplast genome (API) could not be assembled for sks047, and while API contigs were successfully assembled for StAPkA1H1 and sks048 (SI Table 1). API resolved into one and two contigs for sks048 and StAPkA1H1, respectively. Similarly, mitochondrial genome (MIT) contigs were assembled for all three draft assemblies; however, MIT circularisation also failed. Rather than a single sequence, MIT resolved into four, three and one contigs for StAPkA1H1, sks047 and sks048, respectively. All three isolates had reads that span the full-length API and MIT length, though sks047 had <10-fold input read coverage for API, which may have hindered the assembler’s ability to resolve into contigs.

In contrast, API coverage for StAPkA1H1 and sks048 was up to 108x, while MIT coverage for all three isolates was between 292x and 713x. Comparisons with the PKNH reference genome excludes both extranuclear genomes.

### Chromosome structure

Dot plots of draft genome alignment with the PKNH reference shows that the three draft genomes are syntenic with the PKNH reference genome regardless of gaps present in the genomes generated from patient isolates (SI Fig 3). The unplaced sequences in the 00 ‘bin’ chromosomes account for at least 40% of gaps in the three draft genomes (**Error! Reference source not found.**). Indeed, each draft genome’s chromosome structure conforms to that of the PKNH reference genome with uniform coverage across the chromosomes in regions with no gaps (SI Fig 4). This is also apparent in fragmented chromosomes, which retain the same chromosomal structure as PKNH (SI Fig 5). While coverage remains largely uniform, structural variations (>10kb), for example, duplications and inversions, are present in the AM-F assemblies as seen in duplications present in multiple chromosomes in sks047 and sks048 (SI Fig 4B).

Additionally, inversions are present in almost every chromosome, often as inverted duplicate sequences, with the most striking instance observed in chromosome 5 of sks048 (SI Fig 4A iii) where multiple duplicated inversions are observed. Frameshifts, are present across chromosomes in all of the draft genomes (SI Fig 4B). Given the robust clinical isolate draft genome assembly, the frameshifts observed deserve further investigation. Associated gaps do not appear to have impacted the distribution of genes within the draft genomes (Fig 2). Mean annotated gene density shows the PKNH reference genome to have 22.05 genes per 100kbp, StAPkA1H1 to have 18.15, sks047 to have 20.25 and sks048 with 19.80 (Fig 2). Increased gene density may be achieved with manual pseudogene curation since mean gene density is inversely correlated with the number of pseudogenes (Table 2).

**Fig 2.**
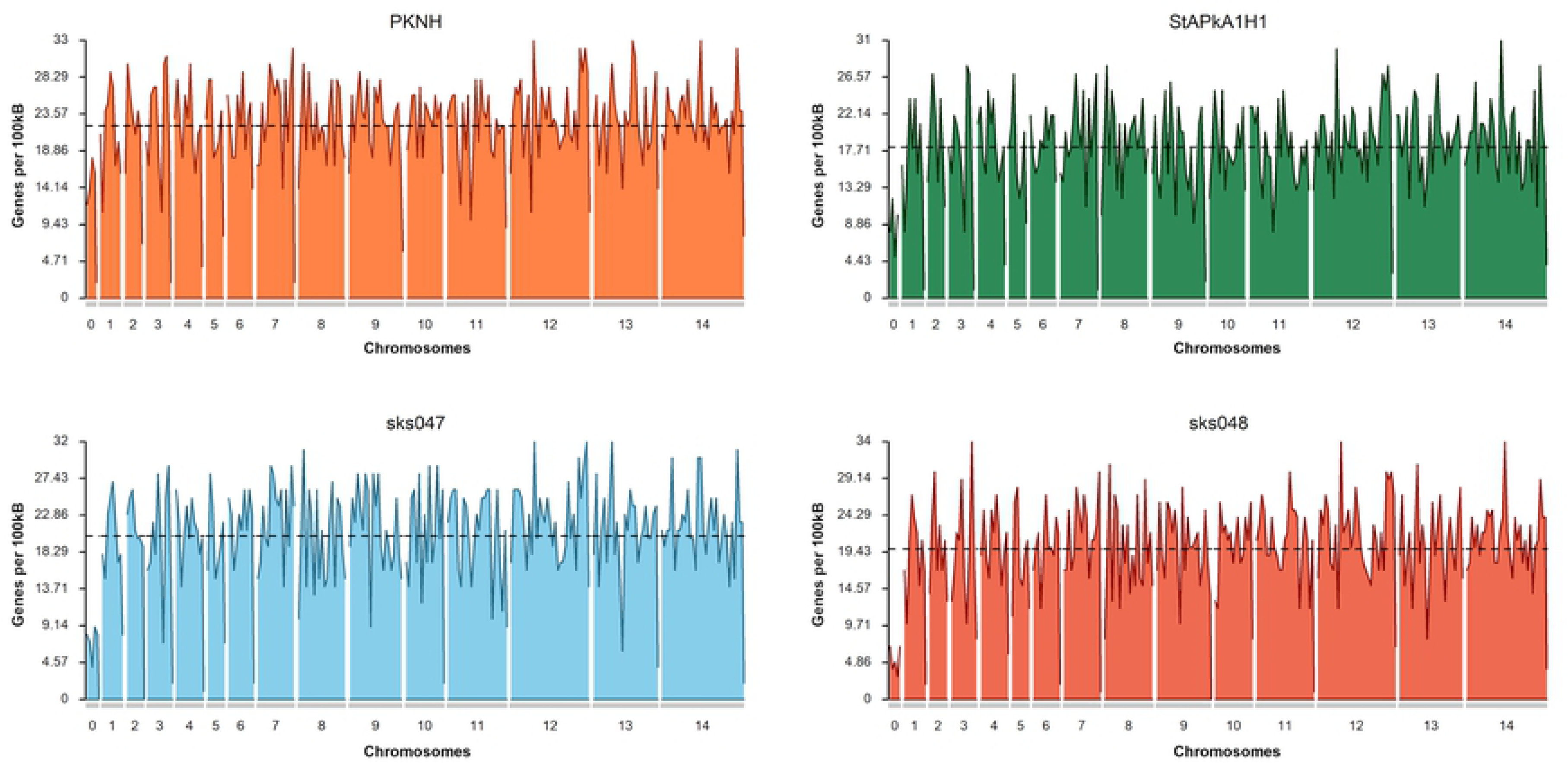
Gene density plots for the *P. knowlesi* PKNH reference genome, StPkA1H1, sks047 and sks048 draft genomes. Gene density is calculated based on the number of identified genes within a sliding window of 100kb. Mean density shows the PKNH reference genome to have 22.05 genes per 100kb, StAPkA1H1 to have 18.15, sks047 to have 20.25 and sks048 with 19.8. Plots were generated using karyoploteR [40].

With the exception of the *SICAvar* Type 1, SICAvar gene fragments and the KIR genes, analysis of the other multigene families reveals similar retention copy number in the three draft genomes and the PKNH reference (Table 3). Given the high similarity between the experimental lines StAPkA1H1 and PKNH in dotplots and other analyses (SI Fig 2A, SI Fig 3A) the expanded number of *KIR* genes in two different laboratory passaged lines, compared with clinical isolates, may reflect gene retention through passive artificial passage. Clinical isolates are effectively wild-type *P. knowlesi* and the lower *KIR* gene copy number in clinical isolates may reflect recombination and selection pressure in mosquito transmission in nature. Chromosomal positional analyses of the *KIR* genes show varied distribution across chromosomes and that only three KIR genes were represented in chromosome 00 in the clinical isolates sks047 and sks048 draft genomes supports the constrained *KIR* gene copy number in nature (SI Fig 6). *SICAvar* genes appear to be distributed across the genome, chromosomes, including the chromosomal extremities with more members annotated than previously reported by Pain et al. (2008), particularly on chromosomes 10, 11 and 12 (SI Fig 7).

**Table 3.** Number of annotated protein copies of the multigene families identified

| *Genes* | *Abbr.* | *PKNH* | *StAPkA1H1* | *sks047* | *sks048* |
| --- | --- | --- | --- | --- | --- |
| *Circumsporozoite protein* | *CSP/CS-TRAP* | 2 | 2 | 2 | 2 |
| *Cytoadherence linked asexual protein/gene* | *CLAG* | 2 | 2 | 2 | 2 |
| *Duffy binding/Duffy-antigen protein [ Erythrocyte binding protein (alpha/beta/gamma)]* | *DBP/DaBP [ ERYBP(a/b/g)]* | 3 | 3 | 3 | 3 |
| *Early transcribed membrane protein* | *ETRAMP* | 9 | 9 | 9 | 9 |
| *Knob-associated histidine-rich protein* | *KAHRP* | 1 | 1 | 1 | 1 |
| *Knowlesi Interspersed Repeats (-like)* | *KIR/KIRL* | 70 | 65 | 28 | 30 |
| *Merozoite surface protein* | *MSP* | 13 | 10 | 10 | 10 |
| *Multidrug resistance (-associated protein)* | *MDRP/MDRaP* | 4 | 3 | 3 | 3 |
| *Reticulocyte binding protein* | *Pknbp/rbp* | 2 | 2 | 2 | 2 |
| *Sporozoite invasion-associated protein* | *SPIAP* | 2 | 2 | 2 | 2 |
| *Tryptophan-rich antigen* | *TrpRA* | 29 | 29 | 30 | 29 |
| *ATP-binding cassette (ABC) transporter* | *ABCtrp* | 15 | 15 | 15 | 15 |
| *Apicomplexan Apetala2 transcription factor* | *ApiAP2* | 29 | 28 | 28 | 28 |
| *Schizont-infected agglutination variant proteins* | *SICAvar* | 109 | 206 | 124 | 160 |

### Structural Variation

Following filtering for length, quality and depth, reads-based structural variants (SVs) were called using the ONT SV pipeline and assembly-based SVs were called using Assemblytics [41]. The reads-based approach returned 1316 and 1398 SVs for sks047 and sks048, respectively (Table 4). The assembly-based approach returned 856 and 839 SVs for sks047 and sks048, respectively (Table 4). The reads-based approach is expected to return more variants due to a higher error rate in the raw reads used compared with the collapsed assembly-based methodology.

**Table 4.** Summary of reads-based and assembly based structural variants

| *Isolate* | *Total SVs* |  | *Insertions* |  | *Deletions* |  |
| --- | --- | --- | --- | --- | --- | --- |
|  | Reads | Assembly | Reads | Assembly | Reads | Assembly |
| *sks047* | 1316 | 856 | 564 | 396 | 752 | 460 |
| *sks048* | 1398 | 839 | 667 | 480 | 731 | 359 |

SVs that exceeded the quality, length and read depth threshold are distributed across the genome on all chromosomes within coding and non-coding regions. Within the 101 shared SVs, 68 were within annotated genes, including within the *SICAvar* and *KIR* multigene families (Supplementary Table 2).

There were different variation signatures between the experimental line StAPkA1H1 compared with the two clinical isolates sks047 and sks048 (Fig 3). StAPkA1H1 had more tandem variants than the clinical isolates, sks047 and sks048. In comparison the clinical isolates show more variation in their repeat sequences with similar insertion and deletion (red and blue) and repeat expansion and contraction (Green and Purple) signatures than StAPkA1H1 (Fig 3).

**Fig 3.**
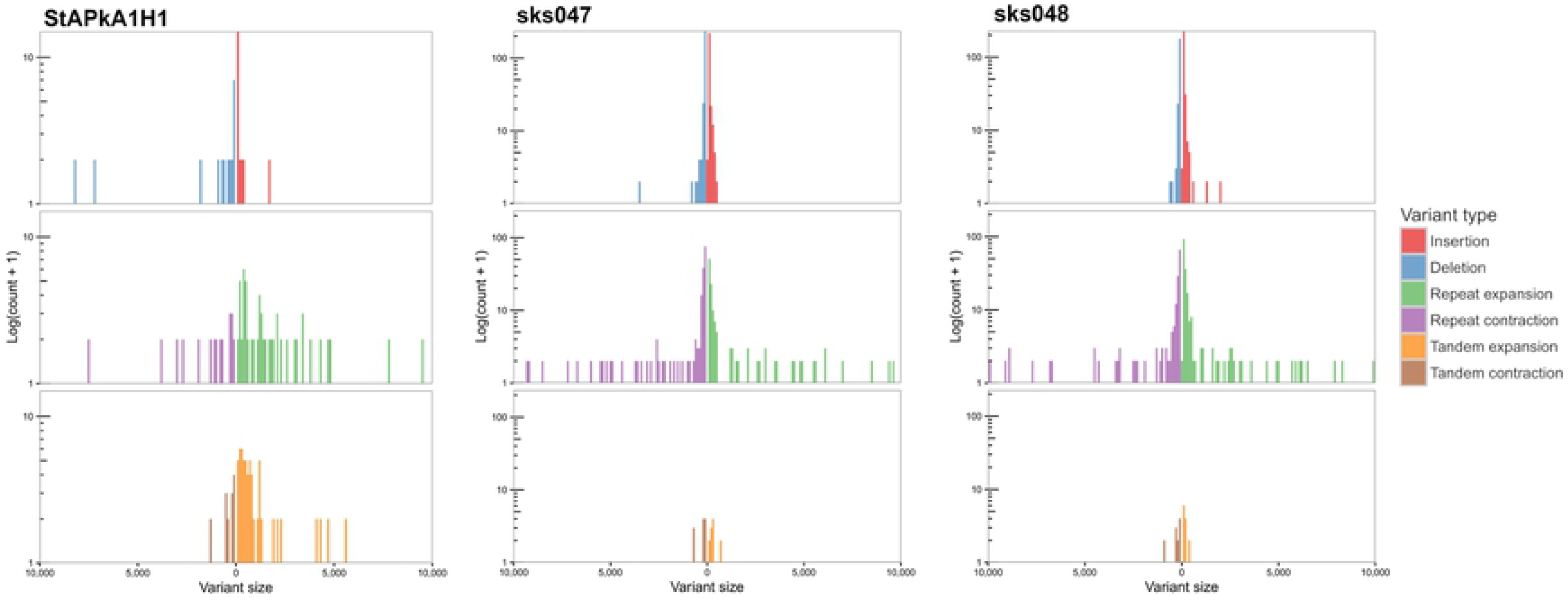
Assembly-based structural variation, size 50 – 10,000bp, of StAPkA1H1, sks047 and sks048 draft genomes against the PKNH reference genome [16]. Nucmer alignment was generated using parameters “—maxmatch -l 100 -c 500” with default and Assemblytics parameters [41]. Expansions (green and orange) refer to insertions that occur within repeat or tandem variants, while contractions (purple and brown) refer to deletions in these regions. More variation is present in the tandem variants (brown and orange) of StAPkA1H1 than those of the draft clinical isolate genomes, sks047 and sks048. In comparison the clinical isolates show more variation in their repeat sequences with similar insertion and deletions (red and blue) and repeat expansion and contraction (green and purple) signatures.

### Gene duplication

Gene duplication was quantified and classified using MCScanX [42]. All genes within the draft genomes for the StAPkA1H1 cultured line and sks047 and sks048 clinical isolates were classified as either: Singleton (no identified duplication; proximal (two identified duplicated genes with <20 genes between them); dispersed (>20 genes between the 2 candidate genes); tandem (duplication events next to each other) and segmental/ whole genome duplication (WGD) (>4co-linear genes with <25 genes between them). To gain an insight into differences in the duplication types we classified duplication types for the *BUSCO* core eucaryotic core control gene population and the *PkSICAvar* type 1, *PkSICAvar* type 2 and the *KIR* multiple gene families of interest in the three draft genomes StAPkA1H1, sks047 and sks048 (Fig 4). The duplication profile of the control population *BUSCO* genes was well matched for each draft genome and also to the *BUSCO* duplication profile for the PKNH reference genome (Mann–Whitney U test StAPkA1H1, *p*=0.92; sks047, *p*=0.67; sks048, *p*=0.66; PKNH *p*= 0.40). Therefore, there was no observed excess duplication types for *BUSCO* genes (Fig 4). However, duplication profiles for the genes annotated *SICAvar* type 1, *SICAvar* type 2 and *KIR* in the draft genomes, StAPkA1H1, sks047 and sks048, were markedly different from the *BUSCO* gene profiles with no evidence for singleton genes (Fig 4). When compared to 100 randomly obtained genes as a population this result profile was statistically significant (Mann–Whitney U test, p < 1.0e-9).

**Fig 4.**
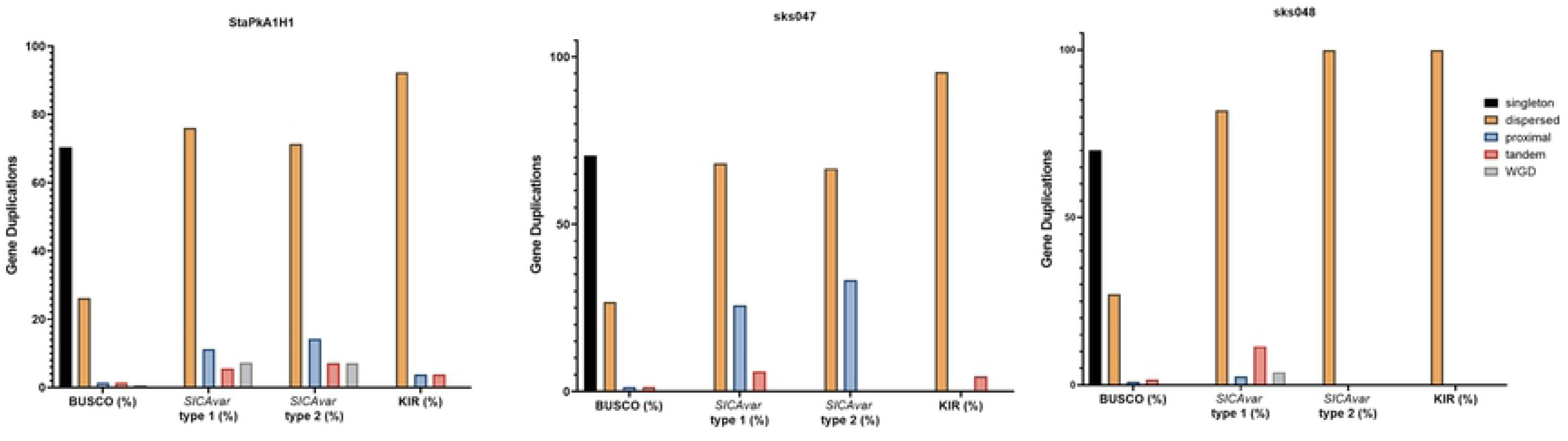
Gene duplication classes for the draft genome assemblies for StAPkA1H1 (experimental line) and the clinical isolates sks047 and sks048. Gene duplication was quantified and classified by MCScanX [42] for all genes in each genome and identified as Singleton (no identified duplication), dark blue bars; proximal (two identified duplicated genes with <20 genes between them), grey bars; dispersed (>20 genes between the 2 candidate genes), orange bars; tandem (duplication events next to each other), yellow bars; and segmental/ whole genome duplication (WGD) (>4co-linear genes with <25 genes between them), light blue bars. The gene pools for each genome were divided into BUSCO (core genome genes) for comparison with the genes making up the *SICAvar* type 1, or *SICAvar* type 2 or *KIR* multiple gene families. The draft genomes, StAPkA1H1, sks047 and sks048, had roughly similar profiles for BUSCO genes. Singletons (blue bars) were absent from the multiple gene families for all of the draft genomes.

### Positive selection: nonsynonymous (dN)/synonymous (dS)substitutions

In order to determine if the *SICAvar* type 1, *SICAvar* type 2 and *KIR* genes are under selection pressure the associated predicted proteins from each genome, StAPkA1H1, PKNH (Reference), sks047 and sks048 were translated into amino acid sequence and clustered into putative orthologous gene clusters containing *SICAvar* type 1, or *SICAvar* type 2 or *KIR* or *BUSCO* (control group) using Orthofinder. The amino acid sequences were aligned and the alignments used to “backtranslate” into nucleotide coding sequences. The mean dN/dS for *SICAvar* type 1, *SICAvar* type 2, *KIR* and BUSCO gene clusters was 2.40, 2.74, 2.35 and 0.35 respectively (Table 5 and SI Fig 8).

**Table 5.** Non-synonymous versus synonymous (dN/dS) analysis of *SICAvar* type 1, *SICAvar* type 2, *KIR* and BUSCO gene clusters represented collectively in the StAPkA1H1, sks047 and sks048 draft genomes and the PKNH Reference genome.

| *Cluster group* | *Cluster count (n)* | *Mean dN/dS per cluster* | *Standard deviation* | *Median* | *Inter quartile range* |
| --- | --- | --- | --- | --- | --- |
| *BUSCO* | 153 | 0.353 | 0.723 | 0.101 | 0.27 |
| *SICAv* type 1 | 15 | 2.4 | 1.31 | 2.37 | 1.86 |
| *SICAv* Type 2 | 5 | 2.74 | 2.54 | 1.83 | 4.02 |
| *KIR* | 26 | 2.35 | 1.19 | 1.99 | 1.5 |

Clusters containing *SICAvar* type 1, or *SICAvar* type 2 or *KIR* genes had a statistically significant greater mean dN/dS value when compared to BUSCO gene clusters (Wilcoxon rank sum test p-value adjustment method Bonferroni: *SICAvar* type 1, = 4.1e-08; *SICAvar* type 2 = 0.0063 and *KIR*, p = 6.7e-13).

### Genomic organisation of suspected ‘weapon’ gene family members

To determine if the gene families of interest: *PkSICAvar* type 1, *PkSICAvar* type 2 and the *KIR* genes, potential virulence or ‘weapon’ genes, are situated in gene sparse regions we quantified the distance from one gene to its neighbour in both a 3 prime (3’) and 5 prime (5’) direction, excluding genes at the start or end of a scaffold. The values were subjected to further analysis using the *BUSCO* results as a negative control (Fig 5A). With the exception of *SICAvar* type 2 in the 3’ direction all ‘weapon’ gene classes had a greater distance to their neighbouring genes in both the 3’ and 5’ direction. In the 3’ direction: Kruskal-Wallis chi-squared = 272.15, df = 4, p-value < 2.2e-16. Wilcoxon signed-rank test, Bonferroni p-value adjustment in comparison to BUSCO: *SICA*var type 1 p = 2e-16, *SICAvar* type 2 *p* = 0.457 and *KIR p* = 1.1e-10. In the 5’ direction all genic distances for the genes of interest were significantly different to the *BUSCO* control population. Kruskal-Wallis chi-squared = 269.33, df = 4, p-value < 2.2e-16. Wilcoxon signed-rank test, Bonferroni p-value adjustment in comparison to BUSCO: *SICAvar* type 1 p = 2e-16; *SICAvar* type 2 p = 0.00123 and *KIR* p = 3.6e-09. Orthofinder gene cluster outputs were further visualised using “UpSets” to determine the membership of genes within each cluster (Fig 5B). The majority of all gene clusters were present in all isolates with the exception of *SICAvar* type 1 gene clusters with 10 – 15 unique *SICAvar* type 1 clusters per isolate. For *KIR* genes, the majority of clusters were shared between all isolates with the exception of a single unique KIR gene cluster in each of sk047 and sk048. The majority of *SICAvar* type 2 genes were orthologues between all isolates with some not identified in sk047 and sk048 (Fig 5B).

**Fig 5.**
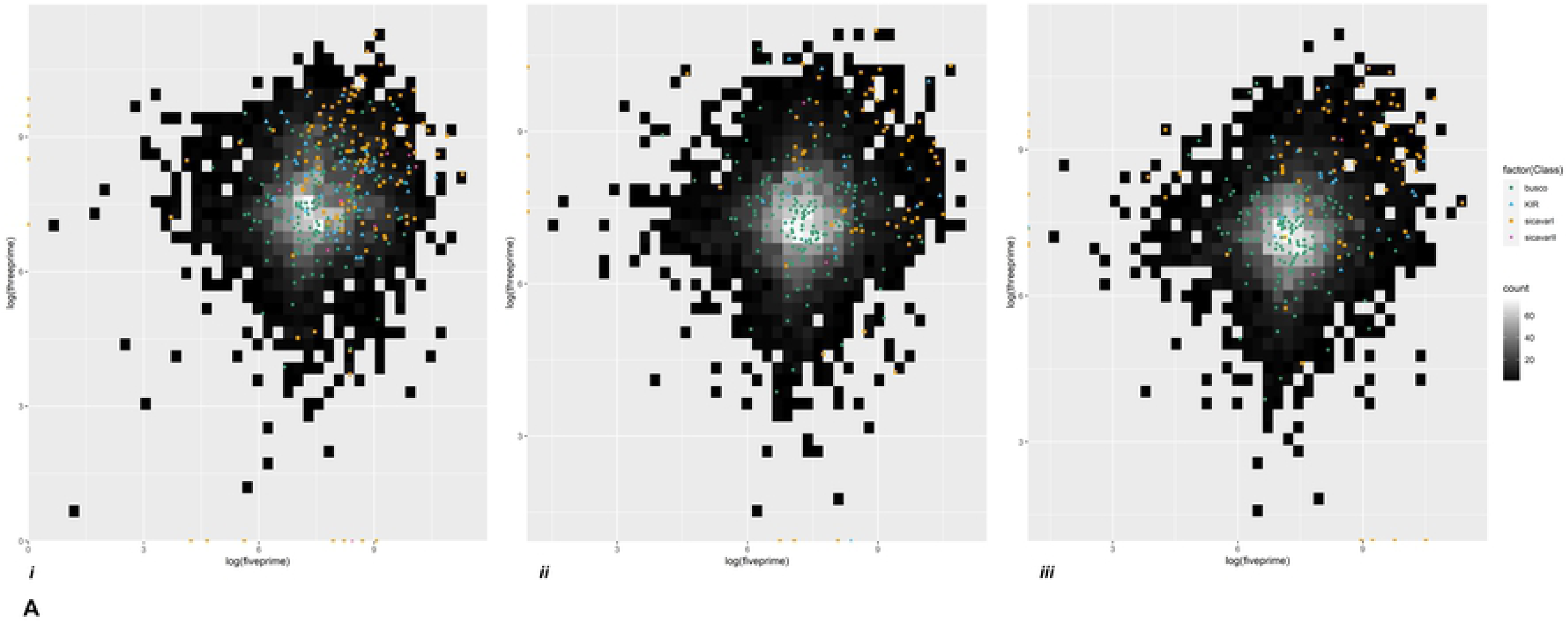
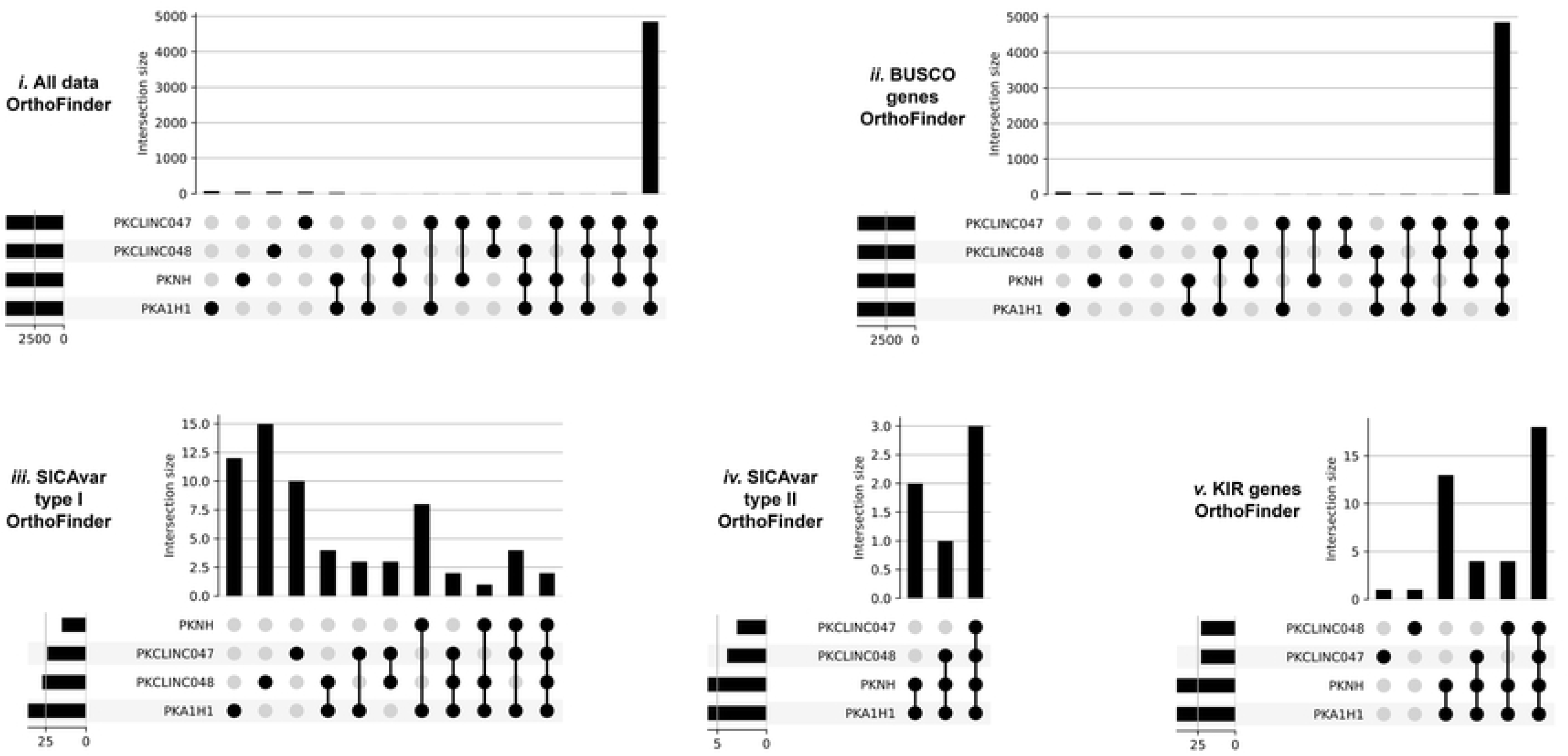
Genomic organisation of ‘weapon’ gene family members. **Fig 5A** Heatmap gene density plots showing 5’ against 3’ intergenic distances (log10) for the draft genomes (i) StAPkA1H1 experimental line, (ii) clinical isolate sk047 and (iii) clinical isolate sk048. Gene density for intergenic distances is represented by colour scale ranging from black (low) to white (high, maximum of 60 genes per bin). Genes classed as BUSCO (green dots), SICAvar type 1 (orange squares), SICAvar type 2 (purple cross) and KIR (blue triangles) are shown. SICAvar type 1 and KIR genes had a significantly greater distance to their neighbouring genes compared with the BUSCO genes (**Wilcoxon signed**-**rank test,** Bonferroni p-value adjustment, p<0.001) suggesting that these gene family members are in gene sparse regions. Genes situated at the start or end of scaffolds were rejected from the analysis. **Fig 5B** Orthofinder gene cluster outputs were visualised using “UpSets” to determine the membership of genes between clusters in the PKNH reference genome [16] and the draft genomes; StAPkA1H1 experimental line, clinical isolate sk047 and clinical isolate sk048. All gene clusters (i), all BUSCO gene clusters (ii), *SICAvar* type 1 gene clusters (iii) *SICAvar* type 2 gene clusters (iv) and *KIR* gene clusters (v) are shown. The majority of all gene clusters were present in all isolates with the exception of *SICAvar* type 1 gene clusters with 10 – 15 *SICAvar* type 1 clusters unique per isolate. For *KIR* genes, the majority of clusters were shared between all isolates with the exception of a single unique KIR gene cluster in each of sk047 and sk048. The majority of *SICAvar* type 2 genes were orthologues between all isolates with some not identified in sk047 and sk048.

## Discussion

Here we present *P. knowlesi* genome sequences assembled from long-read amplification-free sequencing outputs from clinical isolates, essentially wild-type *P. knowlesi*. The new genome sequences are robust and add context to our understanding of *P. knowlesi* genome structure, organisation and variability.

In the first instance we optimized a human leucocyte depletion method to generate high-quality parasite enriched DNA from clinical samples for PCR-free MinION ONT sequencing [45]. To control for our novel methodologies we sequenced the *P. knowlesi* A1-H.1 experimental line and used genome sequence data already available for this line for comparison [34].

Three *Plasmodium knowlesi* genomes were assembled *de novo* from ONT sequence data. Two from *P. knowlesi* clinical isolates (sks047 and sks048) and the other a control genome from the *P. knowlesi* A1-H.1 (StAPkA1H1) experimental line [46]. All three genomes were corrected, polished, and annotated with Racon, Medaka and Companion [47–49]. The clinical isolates sks047 and sks048 were also further corrected with Illumina short-reads using Pilon [50]. Comparison of the *de novo* StAPkA1H1 genome assembled here with the *P. knowlesi* A.1-H1 genome generated using Illumina and PacBio platforms [34] and the *P. knowlesi* reference genome PKNH [16] demonstrated that our sequencing platform and subsequent assembly pipeline produced robust and reliable *de novo P. knowlesi* genome sequences.

ONT long-read sequencing platforms alone can generate *de novo* genomes of good quality [51–53]. However, using our low yield input DNA and in-house pipelines the ONT outputs required correction with high-quality Illumina short reads [17]. Similarly, Lapp et al. generated a reference genome from *P. knowlesi* clone Pk1 using PacBio sequence data that also required additional Hi-C scaffolding [30]. ONT continuously upgrades both their software and hardware and the upgrades are expected to supersede the need for excessive additional correction. For example the recently launched ONT R10 flowcell minimises the inherent homopolymer error rate associated with long-read sequencing technologies [54] and ONT base-calling algorithms report a 32% read error rate reduction [55].

The two clinical isolates (sks047 and sks048) and the control (StAPkA1H1) resolved into 14 chromosomes as expected for *Plasmodium* spp. and one ’bin’ 00 chromosome. The PKNH reference genome also resolves into 14 chromosomes and one bin chromosome where 1.73% of the total sequence comprising 62 genes were assigned [16]. The bin chromosomes (chr00) of StAPkA1H1, sks047 and sks048 contain 1.59%, 2.09% and 1.94% total sequence length with 18, 35 and 25 genes respectively. Sequences placed in chr00 were unable to pass alignment quality thresholds for placement in chromosomes 1-14. For example, when aligned with minimap2 in D-GENIES to produce dotplots, StAPkA1H1 chr00 sequences tended to cluster with PKNH chr00 both representing *P. knowlesi* experimental lines. Failure of sequences to pass quality thresholds would be expected to be randomly distributed genome-wide as observed in sks047 and sks048 chr00 sequences. The observed clustering of StAPkA1H1 chr00 to PKNH chr00 is difficult to explain. It is possible that RagTag may be overriding *‘de novo’* chromosome structuring and ‘forcing’ StAPkA1H1 contigs into a chr00 to fit the pattern set by the PKNH reference genome. Such clustering may be improved by separating the de-chimerisation feature from the scaffolding feature of RagTag and only using the ABACAS feature in Companion to scaffold the contigs. However, both ABACAS and Companion rely on reference-guided chromosome structuring that may also produce similar clustering [37, 48].

During chromosome structuring, we found the minimap2 alignment function of RagTag was unable to resolve chimeric contigs for sks047, sks048 and StAPkA1H1, perhaps, as a function of the algorithm heuristics in minimap2 or localised flaws in our pipeline. Consequently, sections of sks047 chromosomes 02 and 03, which were incorrectly placed in chromosomes 14 and 13 due to chimeric contigs, were successfully corrected using the nucmer aligner function of RagTag.

In general, RagTag struggled to resolve regions of low complexity and high variability, such as telomeric regions. Otto et al., 2018 report that Companion could construct *Plasmodium* chromosomes in their entirety [21]. Indeed, some telomeric sequences were resolved in our *de novo* reference genomes from the clinical isolates, including telomeric sequences identified by Lapp et al. [30]. Furthermore, we report predicted genes within these telomeric regions including some members of the *SICAvar* gene family. More strikingly the *Duffy-binding protein* and *TrpRA* genes are almost exclusively located at the extreme ends of the *de novo* assembled chromosomes.

Our methods were unable to resolve the apicomplast (API) and mitochondrial (MIT) extra-chromosomal genomes completely. In nature, these genomes appear to be circular and possess sequence arrangement that includes a single origin of replication [34]. Our methods may not have disassociated multiple copies of both genomes into single circular API and MIT genomic units. With the exception of the sks047 apicomplast, API and MIT reads were resolved into large contigs with overlapping regions of the same sequence, particularly sks047 MIT.

The genomes interrogated in this study have roughly the same gene - duplication composition as each other, except for the clinical isolate sk047 which did not have any identified segmental whole genome duplications, in contrast to isolate sks048, where 0.28% of genes were identified as segmental. Assembly error can occur when one assembly “over” collapses similar regions, mistaking them for haplotigs, or even outputting excess haplotigs inflating the size or number of segmental duplications. The sk047 and sk048 *de novo* genomes were assembled in exactly the same manner which reduces the probability that this result is an artifact of assembly error. The experimental line, StAPkA1H1, genome had the greatest segmental classed genes (0.63%). It is tempting to speculate that this level of duplication may be the result of many years of less constrained asexual reproduction in tissue culture, reflecting the absence of recombination events during mosquito transmission and vertebrate host-driven selection pressure experienced by wild-type parasites circulating in nature.

We compared the duplication profiles for *SICAvar* type 1, *SICAvar* type 2 and *KIR* gene families, gene families that code for parasite proteins expressed on the surface of infected host red blood cells and that interface with the host, with duplication profiles of the BUSCO genes responsible for normal internal parasite cellular functions. SICAvar type 1, SICAvar type 2 and KIR protein products are antigenically variable and implicated in virulence and are potential “parasite weapons” that require protection from host defence responses. In all of the draft genomes analysed each *SICAvar* type 1*, SICAvar* type 2 and *KIR* gene population had a significantly different duplication profile when compared with 100 randomly selected genes (Mann-Whitney U test: p < 0.001). This suggests that the parasite genome tolerates high levels of duplication at these loci to allow variation, parasite survival and evolution in a hostile host environment. BUSCO core eukaryotic genes are not thought to be under undue selection pressure and were used here as a control gene set to investigate selection pressure. Following gene clustering and dN/dS analysis, clusters which contained *SICAvar* type 1, *SICAvar* type 2 and *KIR* genes had a statistically significantly higher dN/dS values when compared to BUSCO clusters. Non-synonymous substitution over synonymous substitution (dN/dS) values greater than 1.0 are thought to show positive selection pressure. The mean dN/dS for *SICAvar* type 1 gene clusters was 1.31, for *SICAvar* type 2 clusters, 2.54 for *KIR* gene clusters 1.19 while dN/dS for BUSCO gene clusters was 0.35 suggesting that the *SICAvar* type 1*, SICAvar* type 2 and *KIR* gene populations are under strong positive selection pressure. Given that the protein products of these multiple gene family members are expressed at the forefront of parasite host interactions this finding, in addition to multiple copy number within the gene families, would accommodate antigenic variability and makes biological sense by increasing the chance of parasite survival in a hostile host environment.

We then investigated genomic organisation of parasite ‘weapon’ genes to determine if these are located in gene sparse or gene dense regions. With the exception of *SICAvar* type 2 in the 3’ direction, the weapon gene family members had statistically significant greater distances to their neighbouring genes in both the 3’ and 5’ directions compared with BUSCO genes. This suggests that the parasite ‘weapon’ genes are located in gene sparse regions, a genomic arrangement similar to plant pathogens, for example nematodes (Eves van den Akker *et al.,* 2016), aphids (Thorpe *et al.,* 2018), phytophthora (Haas *et al.,* 2009; Thorpe *et al.,* 2021) and fungi (Dong *et al.,* 2015). The ability to tolerate certain genes, *SICAvar* type 1 and *KIR* gene family members in gene sparse, transposon and repetitive rich regions allows the parasite to generate antigenic variability at these important loci while reducing the probability of impacting essential core gene function. The process of genomic regions generating more variation than others is poorly understood, but is termed “the two speed genome” in the field of plant pathogens. In *Plasmodium falciparum*, *Pfemp* 1 gene family members tend to be located in chromosomal sub-telomeric regions. Telomers are unstable with greater rates of recombination in comparison to centromeric regions and this particular location is used to explain the capacity for accruing multiple gene family members and antigenic variability in *P. falciparum* [21].

Following clustering of all genes into their putative orthologous clusters and UpSet visualisation we observed that orthologous versions of *SICAvar* type 1 genes are rarely found in all isolates. With the exception of the PKNH reference genome ([16] where the *SICAvar* gene family members were not well resolved, each of the draft genomes assembled here had between 10 and 15 unique *SICAvar* type 1 gene clusters indicating *SICAvar* type 1 genetic divergence. Indeed, only two *SICAvar* type 1 gene-clusters were shared. The *KIR* genes were less divergent with only one unique gene-cluster in sk048 and in sk047 with most *KIR* gene clusters common between clinical isolates and experimental lines.

The ability to generate variation and maintain fitness is fundamental to the pathogen - host interactions. The pathogen needs to fulfil a successful life span to replicate and disseminate. If the host wins the host pathogen battle, then this marks the end of any particular pathogen germ-line. The ability to generate diversity within the pathogen ‘weapon’ genes increases the chance of pathogen survival. The strong signatures of positive selection pressure and gene duplication on the *P. knowlesi SICAvar* type 1*, SICAvar* type 2 and *KIR* genes irrefutably demonstrate their importance in the fitness and evolution of this particular pathogen.

Here we demonstrate the utility of accessible, portable and affordable PCR-free long-read ONT MinION sequencing to *de novo* assemble *Plasmodium* genomes from very small archived clinical samples. The methods developed provide an opportunity to decrease our reliance on experimental lines to generate data from clinical isolates, in close to real time, and unlock the secrets held in essentially wild-type parasite genomes. The new *P. knowlesi* genomes from clinical isolates presented here provide an important insight into contemporary *P. knowlesi* isolates in Malaysian Borneo and the degree of positive selection exerted, genome wide, on malaria parasites. *P. knowlesi* is a zoonotic infection that is associated with severe and fatal disease and is currently the most prevalent type of malaria causing disease in Malaysia [9]. The *de novo* genomes represent the two dimorphic forms of *P. knowlesi* associated malaria in Malaysian Borneo [17] with some evidence for differential association with disease severity between clusters [35, 56]. On that backdrop the clinically relevant *de novo* genomes will provide an important resource for groups, including ours, reliant on signatures of *P. knowlesi* genome-wide diversity to take forward important research on *P. knowlesi*, from evolutionary biology, zoonotic disease transmission to allelic associations with disease.

## Materials and Methods

### Sample selection

*P. knowlesi* DNA extracted from clinical samples collected with informed consent as part of a non-interventional study were used [35]. The isolates were selected to represent each of the two genetically distinct clusters KH273 (sks047) and KH195 (sks048) of *P. knowlesi* infecting patients in the study cohort [17, 35]. Control *P. knowlesi* DNA was extracted from the experimental line *P. knowlesi* A1-H.1 adapted to *in vitro* culture in human erythrocytes and kindly donated by Robert Moon [46]. In order to distinguish the genome data generated here for *P. knowlesi* A1-H.1 from that already existing we use the unique acronym, StAPkA1H1 [34, 57].

### Plasmodium DNA extraction

Human DNA was depleted from 200 – 400µL thawed clinical samples using a previously described method [45]. Briefly, surviving human leucocytes in thawed samples were removed using anti-human CD45 DynaBeads (ThermoFisher Scientific). The resulting parasite pellet was washed to remove soluble human DNA (hDNA), and parasite enriched DNA (pDNA) was extracted using the QIAamp Blood Mini Kit (QIAGEN) with final elution into 150µL AE Buffer. DNA concentrations were quantified using the Qubit 2.0 fluorometer (Qubit™, Invitrogen) and real-time qPCR on RotorGene (QIAGEN). Recovered DNA was concentrated, and short fragments were removed by mixing 1:1 by volume with AMPureXP magnetic beads (Beckman Coulter) following the manufacturer’s instructions. Briefly, the AMPureXP bead mixture was placed in a magnetic field and DNA bound to the beads was rinsed twice with 70% ethanol before air drying to allow residual ethanol to evaporate. Parasite enriched DNA was eluted in 10uL nuclease-free H_2_O (Ambion). One µl of recovered DNA concentrate was used for DNA quantification using Qubit Fluorimetry (ThermoFisher Scientific) and 7.5µl taken forward for sequencing library preparation.

### Library preparation and Sequencing

Parasite enriched DNA was sequenced using the Oxford Nanopore Technologies (ONT) MinION long-read sequencing platform. Library preparations were selected to suit PCR-free sequencing for the small pDNA quantities available to study (∼400ng). Sequencing libraries were prepared following the manufacturer’s instructions for the SQK-RBK004 ONT sequencing kit. Sequencing was performed using R9.4.1 flowcells or R10 flowcells [45]. Previously sequenced Illumina reads for the patient isolates (sks047 and sks048) were retrieved from the European Nucleotide Archive, with accession codes ERR366425 and ERR274221, respectively [17]. Further short-read sequencing was carried out on PCR-enriched DNA using the Illumina MiSeq platform at the London School of Hygiene and Tropical Medicine and methods established by Diez Benavente et al., [58].

### Reference Genomes

For chromosome scaffolding and quality assessment comparison, the *P. knowlesi* PKNH reference genome [16] (version 2) was downloaded from Sanger (ftp://ftp.sanger.ac.uk/pub/genedb/releases/latest/Pknowlesi/#). In addition, further comparisons were carried out using the *P. knowlesi* PkA1H1 reference genome [34]from NCBI [accession code: GCA_900162085].

### De novo genome Assembly

MinION FAST5 file outputs were locally base called using the high accuracy model of the guppy basecaller (v4.0.15; Ubuntu 19.10; GTX1060) with the following parameters: *′-r -v -q 0 --qscore-filtering -x auto*′. Demultiplexing was carried out using qcat software (v1.1.0) with the ′*--detect-middle --trim -k --guppy*′ parameters, then adapter removal with porechop (v0.2.4) using default parameters and the most recent aversions released from ONT technologies. Human DNA (hDNA) contamination was removed from the adapter-free reads by alignment against the human GRCh38.p13 reference genome (retrieved from NCBI accession code: GCF_000001405.39) [59] using minimap2 (v2.17); [60] with ′*-ax map-ont*′ default parameters. Unmapped reads were separated from the binary sequence alignment (BAM) file with samtools (v1.10; [61, 62] and converted back to FASTQ by bedtools (v2.29.2) [63] for *de novo* genome assembly using Flye, (v2.8.1) [64] with an expected genome size of 25Mb and ′*--nano-raw′* default parameters. Successful assemblies were assessed for contamination using BlobTools (v1.0.1) [65]. Contigs not taxonomically assigned as Apicomplexan were discarded.

### Assembly Polishing and Correction

Draft assemblies were polished using four iterations of racon (v1.4.13) [49]; in the default setting retaining raw long-read isolate sequence reads which did not align to the human GRCh38.p13 (henceforth parasite-reads). As part of the polishing step, alignments of parasite-reads against the draft assembly were performed with minimap2 (v2.17; [60] . A consensus sequence was subsequently generated from the racon output using medaka (v1.0.3; default settings) [47]. Further polishing and correction was carried out using Illumina paired-end reads where available, using three iterations of pilon (v1.23) default parameters with ′*-Xmx120G, --tracks, --fix all, circles*′) [50].

### Masking repetitive elements

The *P. knowlesi* PKNH reference mitochondrial (MIT) and apicoplast (API) sequences were extracted and individually aligned against draft *P. knowlesi* assemblies using MegaBLAST (v.2.9; default parameters) [16, 66]. Contigs which aligned to the reference PKNH MIT and API genomes were subsequently removed and circularised on Circlator (v1.5.5) [67] with the command ′*circlator all -- data_type nanopore-raw --bwa_opts "-x ont2d" --merge_min_id 85 --merge_breaklen 1000*′.

API/MIT-free draft assemblies (henceforth AM-F assemblies) were taken forward through RepeatModeler (v1.0.10) [68]and the outputs utilised as input for Censor [69] where the options ′*Eukaryota*′ and ′*Report simple repeats*′ were selected. Identified transposable elements and repeats in the censor outputs were classified based on the class of repeats to make a repeat library for each AM-F assembly. Repeat libraries of each AM-F assembly were combined and misplaced, redundant, sequences removed with CD-HIT (v4.8.1; ′*-c 1.0 -n 10 -d 0 -g 1 -M 60000*′ parameters) [70, 71]. This generates a singular ’master’ repeat library encompassing the non-redundant list of identified elements across the three AM-F assemblies.

With the master repeat library, RepeatMasker (v4.0.7) was run on each AM-F assembly producing a tab-separated value (TSV) output of the identified repeats in the assembly. Then, using ’One Code to Find Them All’ (OCFTA) [72], each TSV file was parsed to clarify further repeat positions found by RepeatMasker. Next, the LTRHarvest [73] module of GenomeTools (v1.6.1) [74] was used to find secondary structures of long terminal repeats (LTRs) and other alternatives in the AM-F assemblies. Here, the ’*suffixerator*’ function was implemented with ′*-tis -suf -lcp -des -ssp -sds -dna*′ parameters while the ’*ltrharvest*’ function was run with ′*-mintsd 5 -maxtsd 100*’ parameters. Concurrently, TransposonPSI was also used on the AM-F assemblies with default parameters to find repeat elements based on their coding sequences.

Redundant repeat element sequences were removed from the outputs of RepeatMasker, OCTFA, LTRHarvest and TransposonPSI using a custom script, to generate a genome feature file (GFF3) where each transposable and repetitive element of each AM-F assembly is represented once. Then, within each draft assembly, repeat elements were masked using the coordinates present in the non-redundant GFF3 file and the ’*maskfasta*’ function of bedtools (v2.27; default settings and *′-soft*′).

### Prediction and Annotation

The masked AM-F assemblies were checked for chimeric contigs using Ragtag (v1.0.1) [75] where both the ’*correct*’ and ’*scaffold*’ functions were run with the ′*--debug --aligner nucmer --nucmer-params=’-maxmatch -l 100 -c 500’′* parameters [61, 62].

With the chimeric contigs broken, masked AM-F assemblies were uploaded on the Companion webserver [48] for gene prediction and annotation using the sequence prefix of ’PKA1H1_STAND’ for the cultured experimental line (StAPKA1H1) and ’PKCLINC’ for patient isolates (sks047 and sks048). Companion software was run with no transcript evidence, 500bp minimum match length and 80% match similarity for contig placement, 0.8 AUGUSTUS [76] score threshold and taxid 5851.

Additionally, pseudochromosomes were contiguated, reference proteins were aligned to the target sequence, pseudogene detection was carried out, and RATT was used for reference gene models. *Comparative Genomics, Quality Assessment and Analyses*

As the pipeline progressed, assembly metrics were checked with assembly-stats (v1.0.1) and pomoxis (v0.3.4). Additionally, draft genomes were further assessed for completeness and accuracy using BUSCO(v5.0) with *′-l plasmodium_odb10 -f -m geno --long*′ parameters [77]. GFF3 files generated on Companion were parsed for genes of interest, including multigene families known to span the core genome and telomeric regions. Chromosomes of the annotated AM-F draft genomes were individually aligned against the corresponding *P. knowlesi* PKNH reference chromosome [16] with minimap2 parameters ′*-ax asm5*′. Resulting alignment files were analysed on Qualimap (v.2.2.2) [78] with parameters ′ *-nw 800 -hm 7*′. Gene density, chromosome structure and multigene family plots were generated using the karyoploteR visualisation package [40]. Dotplots to identify repetitions, breaks and inversions were generated from minimap2 whole genome alignments using D-GENIES default settings [79].

### Structural Variant Analyses

The StAPkA1H1 draft genome, assembled here, was used as the reference for structural variant calling and subsequent variant annotation to ensure parity across sequencing technologies. Read alignment-based structural variant calling (henceforth reads-based) was achieved using the Oxford Nanopore structural variation pipeline (ONTSVP) (https://github.com/nanoporetech/pipeline-structural-variation) while the assembly-based approach was completed with Assemblytics [41].

Using a modified Snakefile, FASTQ isolate parasite-reads and the StAPkA1H1 draft genome; the ONTSVP first parses the input reads using catfishq (https://github.com/philres/catfishq) and seqtk (https://github.com/lh3/seqtk) before carrying out alignment using lra with parameters ′*-ONT -p s′* [80]. The resulting alignment file was sorted and indexed with samtools and read coverage was then calculated using mosdepth (′*-x -n -b 1000000*′) [81]. Structural variants (SV) were called by cuteSV [82] with parameters ′*--min-size 30 --max-size 100000 --retain_work_dir --report_readid -- min_support 2*′. Variants were subsequently filtered for length, depth, quality, and structural variant type (SVTYPE) such as insertions (INS) by default, before filtered variants were sorted and indexed. Failed SV types were manually filtered based on length and quality alone to determine the presence of high-quality, low-occurrence variants.

For the assembly-based structural variant calling for the clinical isolates sks047 and sks048 and StAPkA1H1 draft genomes were aligned against the PKNH reference genome [16] using nucmer with ′--maxmatch *-l 100 -c 500*′ parameters and outputs uploaded onto Assemblytics (http://assemblytics.com) [41] with default parameters and a minimum SV length of 30bp. BEDfile outputs of Assemblytics were converted to variant call format (VCF) file using SURVIVOR (v1.0.7) [83]. VCF files for successful reads-based and assembly-based SV calling as well as the failed SV-type VCF files were further filtered to remove any variants less than 50bp in length and less than Q5 in quality using a bcftools one-liner (https://github.com/samtools/BCFtools). A quality filter was not applicable for the assembly-based approach due to the lack of quality information in the original BEDfile output of Assemblytics. Variants exceeding these thresholds were annotated with vcfanno (v0.3.2) [41] and subsequently sorted and indexed. Annotated variants, relevant BAM alignment files and GFF files were visualised on IGV [84]. Using IGV, a gene locus previously identified to be associated with dimorphism – *PknbpXa* [17]—was analysed to determine the presence of structural variants. Summary statistics were calculated using the ’stats’ function of SURVIVOR with parameters ′-1 -1 -1′. VCF files were compared using the ‘isec’ function of bcftools with default settings, including analyses of the variants present within genes.

### Duplication, clustering, genomic organisation and dN/dS analyses

Scripts used can be found here: https://github.com/peterthorpe5/plasmidium_genomes. Gene duplication analyses were performed using the similarity searches from DIAMOND-BlastP (1e-5) with MCSanX toolkit[42]. Orthologues clustering and dN/dS was performed as described in [43]. Briefly, Orthofinder (v2.2.7) [85] was used to cluster all the amino acids sequences for the genomes used in this study. The resulting sequences from the clusters of interest were aligned using MUSCLE (v3.8.1551) [86] and refined using MUSCLE. The resulting amino acid alignment was used as a template to back-translate the nucleotide coding sequence using Biopython for subsequent nucleotide alignment [87]. The nucleotide alignment was filtered to remove any insertions and deletions and return an alignment with no gaps using trimAL (v1.4.1)[88]. The resulting alignment was subjected to dN/dS analysis using Codonphyml (v1.00 201407.24) (-m GY --fmodel F3X4 -t e -f empirical -w g -a e ) [44]. Genomic organisation of classes of genes of interest was performed as described in [43, 89, 90]. For UpSet visualization the scripts can be found in the github link above.

## Author Contributions

*DRO, data curation, formal analyses, investigation, methodology, visualization; PT, formal analyses, software, investigation, supervision; EDB, Supervision; SC, resources; FM, Resources; RM, Resources, editing; TGC, supervision, draft editing; JCS, conceptualization, funding acquisition, methodology, project administration, resources, supervision, writing preparing draft*.

## Acknowledgements

We would like to acknowledge Dr Joseph Ward for help with software and resources and Dr Fiona Cook for providing resources for optimising methodologies.

## Funding

DRO is supported by the Wellcome Trust ISSF award 204821/Z/16/Z. Bioinformatics and computational biology analyses were supported by the University of St Andrews Bioinformatics Unit (AMD3BIOINF), funded by Wellcome Trust ISSF award 105621/Z/14/Z and 204821/Z/16/Z. The sample BioBank was compiled with informed consent (Medial Research Council, www.mrc.ac.uk, grant G0801971). Genome sequencing was supported by Tenovus Scotland (T16/03). TGC is funded by the Medical Research Council UK (Grant no. MR/M01360X/1, MR/N010469/1, MR/R025576/1, and MR/R020973/1) and BBSRC (Grant no. BB/R013063/1). SC is funded by Medical Research Council UK grants (ref. MR/M01360X/1, MR/R025576/1, and MR/R020973/1).

## Data availability

Genomes are in the process of being deposited in NCBI. Raw reads associated with these genomes are also deposited under the same accession number. Scripts used to generate the data in this project are available in github: https://github.com/damioresegun/Pknowlesi_denovo_genome_assembly and https://github.com/peterthorpe5/plasmidium_genomes.

## Supplementary Information (SI)

**SI File**

Full assembly statistic metrics for the PKNH reference sequence with the apicoplast and mitochondrial sequences included (and excluded: PKNH_noAPI/MIT), Cultured PkA1H1 isolate (StAPkA1H1), sks047 and sks048. The file contains metrics amalgamated from the outputs of Companion, AGAT, QUAST, BUSCO and Assembly-stats. In addition, specific features of the genomes have been separated into sub-pages, such as tRNAs and rRNAs.

**SI Figures**

**SI Fig 1.** Whole genome coverage across chromosomes of the StAPkA1H1, sks047 and sks048 draft genomes against the PKNH reference genome [16]. Coverage and plots generated using Qualimap are shown. The red trace shows troughs that indicate regions of low coverage. Coverage appears more stable in StAPkA1H1 (i) than in the clinical isolates sks047 (ii) and sks048 (iii) indicating higher variability in the contemporary *P. knowlesi* genomes than in the experimental line when compared with the reference.

**SI Fig 2.** Alignments of chromosome 00 (bin) for StAPkA1H1, sks047 and sks048 against the whole PKNH genome [16]. Minimap2 alignments of the bin chromosomes against the entire PKNH reference genome with a 1kbp alignment length filter. The ‘bin’ chromosomes contain sequence fragments that could not be confidently resolved into a particular chromosome during the scaffolding process. StAPkA1H1(i) shows a concentration of sequences aligned to the PKNH ‘bin’ chromosome 00 (green box), while no clustering is evident in sks047(ii) and sks048 (iii).

**SI Fig 3.** Whole-genome alignment of StAPkA1H1, sks047 and sks048 against the *P. knowlesi* PKNH reference genome [16]. Dotplots to identify repetitions, breaks and inversions were generated from minimap2 whole draft genome alignments for StAPkA1H1(i), sks047 (ii) and sks048 (iii) using D-GENIES default settings [79].

The PKNH chromosomes 00 – 14 are shown on the x-axes at the top and size given on the bottom in MB. Draft genome chromosomes 00 – 14 are shown on the right y-axes and size in MB on the left. The line indicates gene synteny between each draft and the PKNH reference genome. Red boxes show where the draft ‘00’ chromosomes align with PKNH chromosome ‘00’.

**SI Fig 4.** Dot plots showing draft genomes aligned against the PKNH reference genome [16] with minimum alignment 10kB. SI Fig 4A Chromosome 5 is given for StAPkA1H1 (i), sks047 (ii) and sks048 (iii), as an example where frameshifts are outlined in purple, gaps outlined in orange, inversions outlined in green and inverted repeats in red. Duplications are not shown. SI Fig 4B Shows dot plots of alignments of all chromosomes for StAPkA1H1 (i), sks047 (ii) and sks048 (iii) plotted against the PKNH reference genome [16] with minimum alignment 10kB. Gaps, frameshifts and large structural variants are dispersed across the draft genomes are shown.

**SI Fig 5.** Mauve plot of chromosome 08 for StAPkA1H1, sks047, sks048 and the PKNH [16] reference genome. Chromosome 8 of the PKNH reference shows more fragmentation than other chromosomes in the genome which may have influenced the chromosome structure inferred for the

draft genomes generated here. Extensive mosaicism has been described in *P. knowlesi* chromosome 8 due to an overrepresentation of genes expressed in the mosquito stage of the parasite’s life cycle [57]. Regions of low coverage are still apparent in the draft genomes compared with the PKNH reference genome (red boxes).

**SI Fig 6.** Positioning of non-*SICAvar* multigene family members are shown for the PKNH reference genome and the three draft genomes using karyoploteR [40].

**SI Fig 6A** *P. knowlesi* PKNH(Pain et al 2008);
**SI Fig 6B** StAPkA1H1 experimental line;
**SI Fig 6C** clinical isolate sks047;
**SI Fig 6D** clinical isolate sks048.

Genes are shown as black squares marked along the chromosome linear map. Genes on the positive strand appear above the map line and those on the negative strand below. Identified members of select multigene families are given and colour coded based on being on the positive or negative strand e.g *TrpRA* on the positive strand is slate blue and coral on the negative strand. The *SICAvar* gene family members are presented separately in **SI Fig 7**.

**SI Fig 7.** Positioning of *SICAvar* multigene family members are shown for the PKNH reference genome and the three draft genomes using karyoploteR [40].

***SI Fig 7A*** *P. knowlesi* PKNH(Pain et al 2008);
***SI Fig 7B*** StAPkA1H1 experimental line;
***SI Fig 7C*** clinical isolate sks047;
***SI Fig 7D*** clinical isolate sks04*8*.

Annotated genes are shown as black squares marked along the chromosome linear map.

Genes on the positive strand appear above the map line and those on the negative strand below. *SICAvar* genes and gene fragments on the positive strand are in red font and on the negative strand in green font.

**SI Fig 8.** Box plot to represent dN/dS ratios for gene-clusters from each gene type*: BUSCO; KIR; SICAvar type 1* and *SICAvar type 2* in the combined dataset from draft genomes StAPkA1H1, sks047, sks048 and the PKNH reference [16]. There were 154 BUSCO, 27 KIR, *15* SICAvar *type 1* and *5* SICAvar type 2 gene clusters with mean number of genes 5, 8.59, 32.2 and 18.2 per cluster respectively. Clusters containing *SICAvar* type 1, or *SICAvar* type 2 or *KIR* genes had a statistically significant greater mean dN/dS value when compared to BUSCO gene clusters (Wilcoxon rank sum test p-value adjustment method Bonferroni: *SICAvar* type 1, = 4.1e-08; *SICAvar* type 2 = 0.0063 and *KIR*, p = 6.7e-13) suggesting these gene family members are under selection pressure.

**SI Tables**

**SI Table 1.** Length of non-nuclear DNA content present in the *P. knowlesi* PKNH [16] and PkA1H1 [34] reference in comparison to the three generated draft genomes.

**SI Table 2.** Comparisons of variant call format (VCF) files of sks047 and sks048 against StAPkA1H1 draft genome.

Legend to supplementary Table 2. Legend to Supplementary Table 2: Comparisons were achieved after analysis using the intersect (isec) function of bedtools. Assembly-based SV calling approach utilised Assemblytics [41] to call variants between the isolate draft genomes sks047, sks048 and the StAPkA1H1 draft genome. Reads-based SV calling approach used input reads of the isolate draft genomes against the StAPkA1H1draft genome to call variants with the Oxford Nanopore Structural Variation pipeline.

## Notes

### Competing Interest Statement

The authors have declared no competing interest.

## References

1. Knowles R, Gupta BMD. A Study of Monkey-Malaria, and Its Experimental Transmission to Man. Ind Med Gaz. 1932;67(6):301–20. Epub 1932/06/01. PubMed PMID: 29010910; PubMed Central PMCID: PMCPMC5231565.

2. Butcher GA, Mitchell GH. The role of Plasmodium knowlesi in the history of malaria research. Parasitology. 2018;145(1):6–17. Epub 2016/11/11. doi: 10.1017/S0031182016001888. PubMed PMID: 27829470.

3. Galinski MR, Lapp SA, Peterson MS, Ay F, Joyner CJ, KG LER, et al. Plasmodium knowlesi: a superb in vivo nonhuman primate model of antigenic variation in malaria. Parasitology. 2018;145(1):85–100. Epub 2017/07/18. doi: 10.1017/S0031182017001135. PubMed PMID: 28712361; PubMed Central PMCID: PMCPMC5798396.

4. Pasini EM, Zeeman AM, Voorberg VANDERWA, Kocken CHM. Plasmodium knowlesi: a relevant, versatile experimental malaria model. Parasitology. 2018;145(1):56–70. Epub 2016/12/13. doi: 10.1017/S0031182016002286. PubMed PMID: 27938428.

5. Chin W, Contacos PG, Coatney GR, Kimball HR. A Naturally Acquited Quotidian-Type Malaria in Man Transferable to Monkeys. Science. 1965;149(3686):865. Epub 1965/08/20. doi: 10.1126/science.149.3686.865. PubMed PMID: 14332847.

6. Chin W, Contacos PG, Collins WE, Jeter MH, Alpert E. Experimental mosquito-transmission of Plasmodium knowlesi to man and monkey. Am J Trop Med Hyg. 1968;17(3):355–8. Epub 1968/05/01. doi: 10.4269/ajtmh.1968.17.355. PubMed PMID: 4385130.

7. Singh B, Kim Sung L, Matusop A, Radhakrishnan A, Shamsul SS, Cox-Singh J, et al. A large focus of naturally acquired Plasmodium knowlesi infections in human beings. Lancet. 2004;363(9414):1017–24. Epub 2004/03/31. doi: 10.1016/S0140-6736(04)15836-4. PubMed PMID: 15051281.

8. World-Health-Organization. World Malaria Report 2019. Geneva: 2019 Licence: CC BY-NC-SA 3.0 IGO.

9. Chin AZ, Maluda MCM, Jelip J, Jeffree MSB, Culleton R, Ahmed K. Malaria elimination in Malaysia and the rising threat of Plasmodium knowlesi. J Physiol Anthropol. 2020;39(1):36. Epub 2020/11/25. doi: 10.1186/s40101-020-00247-5. PubMed PMID: 33228775; PubMed Central PMCID: PMCPMC7686722.

10. Cox-Singh J, Davis TM, Lee KS, Shamsul SS, Matusop A, Ratnam S, et al. Plasmodium knowlesi malaria in humans is widely distributed and potentially life threatening. Clin Infect Dis. 2008;46(2):165–71. Epub 2008/01/04. doi: 10.1086/524888. PubMed PMID: 18171245; PubMed Central PMCID: PMCPMC2533694.

11. Cox-Singh J, Hiu J, Lucas SB, Divis PC, Zulkarnaen M, Chandran P, et al. Severe malaria - a case of fatal Plasmodium knowlesi infection with post-mortem findings: a case report. Malar J. 2010;9:10. Epub 2010/01/13. doi: 10.1186/1475-2875-9-10. PubMed PMID: 20064229; PubMed Central PMCID: PMCPMC2818646.

12. Daneshvar C, Davis TM, Cox-Singh J, Rafa’ee MZ, Zakaria SK, Divis PC, et al. Clinical and laboratory features of human Plasmodium knowlesi infection. Clin Infect Dis. 2009;49(6):852–60. Epub 2009/07/29. doi: 10.1086/605439. PubMed PMID: 19635025; PubMed Central PMCID: PMCPMC2843824.

13. Daneshvar C, William T, Davis TME. Clinical features and management of Plasmodium knowlesi infections in humans. Parasitology. 2018;145(1):18–31. Epub 2017/01/27. doi: 10.1017/S0031182016002638. PubMed PMID: 28122651.

14. Cox-Singh J. Plasmodium knowlesi: experimental model, zoonotic pathogen and golden opportunity? Parasitology. 2018;145(1):1–5. Epub 2017/11/17. doi: 10.1017/S0031182017001858. PubMed PMID: 29144211.

15. Cox-Singh J, Culleton R. Plasmodium knowlesi: from severe zoonosis to animal model. Trends Parasitol. 2015;31(6):232–8. Epub 2015/04/04. doi: 10.1016/j.pt.2015.03.003. PubMed PMID: 25837310.

16. Pain A, Bohme U, Berry AE, Mungall K, Finn RD, Jackson AP, et al. The genome of the simian and human malaria parasite Plasmodium knowlesi. Nature. 2008;455(7214):799–803. Epub 2008/10/10. doi: 10.1038/nature07306. PubMed PMID: 18843368; PubMed Central PMCID: PMCPMC2656934.

17. Pinheiro MM, Ahmed MA, Millar SB, Sanderson T, Otto TD, Lu WC, et al. Plasmodium knowlesi genome sequences from clinical isolates reveal extensive genomic dimorphism. PLoS One. 2015;10(4):e0121303. Epub 2015/04/02. doi: 10.1371/journal.pone.0121303. PubMed PMID: 25830531; PubMed Central PMCID: PMCPMC4382175.

18. Harrison TE, Reid AJ, Cunningham D, Langhorne J, Higgins MK. Structure of the Plasmodium-interspersed repeat proteins of the malaria parasite. Proc Natl Acad Sci U S A. 2020;117(50):32098–104. Epub 2020/12/02. doi: 10.1073/pnas.2016775117. PubMed PMID: 33257570; PubMed Central PMCID: PMCPMC7749308.

19. Wahlgren M, Goel S, Akhouri RR. Variant surface antigens of Plasmodium falciparum and their roles in severe malaria. Nat Rev Microbiol. 2017;15(8):479–91. Epub 2017/06/13. doi: 10.1038/nrmicro.2017.47. PubMed PMID: 28603279.

20. Gardner MJ, Hall N, Fung E, White O, Berriman M, Hyman RW, et al. Genome sequence of the human malaria parasite Plasmodium falciparum. Nature. 2002;419(6906):498–511. Epub 2002/10/09. doi: 10.1038/nature01097. PubMed PMID: 12368864; PubMed Central PMCID: PMCPMC3836256.

21. Otto TD, Bohme U, Sanders M, Reid A, Bruske EI, Duffy CW, et al. Long read assemblies of geographically dispersed Plasmodium falciparum isolates reveal highly structured subtelomeres. Wellcome Open Res. 2018;3:52. Epub 2018/06/05. doi: 10.12688/wellcomeopenres.14571.1. PubMed PMID: 29862326; PubMed Central PMCID: PMCPMC5964635.

22. Abdi AI, Hodgson SH, Muthui MK, Kivisi CA, Kamuyu G, Kimani D, et al. Plasmodium falciparum malaria parasite var gene expression is modified by host antibodies: longitudinal evidence from controlled infections of Kenyan adults with varying natural exposure. BMC Infect Dis. 2017;17(1):585. Epub 2017/08/25. doi: 10.1186/s12879-017-2686-0. PubMed PMID: 28835215; PubMed Central PMCID: PMCPMC5569527.

23. Andrade CM, Fleckenstein H, Thomson-Luque R, Doumbo S, Lima NF, Anderson C, et al. Increased circulation time of Plasmodium falciparum underlies persistent asymptomatic infection in the dry season. Nat Med. 2020;26(12):1929–40. Epub 2020/10/28. doi: 10.1038/s41591-020-1084-0. PubMed PMID: 33106664.

24. Hviid L, Jensen AT. PfEMP1 - A Parasite Protein Family of Key Importance in Plasmodium falciparum Malaria Immunity and Pathogenesis. Adv Parasitol. 2015;88:51–84. Epub 2015/04/26. doi: 10.1016/bs.apar.2015.02.004. PubMed PMID: 25911365.

25. Jensen AR, Adams Y, Hviid L. Cerebral Plasmodium falciparum malaria: The role of PfEMP1 in its pathogenesis and immunity, and PfEMP1-based vaccines to prevent it. Immunol Rev. 2020;293(1):230–52. Epub 2019/09/29. doi: 10.1111/imr.12807. PubMed PMID: 31562653; PubMed Central PMCID: PMCPMC6972667.

26. Lavstsen T, Turner L, Saguti F, Magistrado P, Rask TS, Jespersen JS, et al. Plasmodium falciparum erythrocyte membrane protein 1 domain cassettes 8 and 13 are associated with severe malaria in children. Proc Natl Acad Sci U S A. 2012;109(26):E1791-800. Epub 2012/05/24. doi: 10.1073/pnas.1120455109. PubMed PMID: 22619319; PubMed Central PMCID: PMCPMC3387094.

27. Milner DA, Jr. Malaria Pathogenesis. Cold Spring Harb Perspect Med. 2018;8(1). Epub 2017/05/24. doi: 10.1101/cshperspect.a025569. PubMed PMID: 28533315; PubMed Central PMCID: PMCPMC5749143.

28. Shabani E, Hanisch B, Opoka RO, Lavstsen T, John CC. Plasmodium falciparum EPCR-binding PfEMP1 expression increases with malaria disease severity and is elevated in retinopathy negative cerebral malaria. BMC Med. 2017;15(1):183. Epub 2017/10/14. doi: 10.1186/s12916-017-0945-y. PubMed PMID: 29025399; PubMed Central PMCID: PMCPMC5639490.

29. Tessema SK, Nakajima R, Jasinskas A, Monk SL, Lekieffre L, Lin E, et al. Protective Immunity against Severe Malaria in Children Is Associated with a Limited Repertoire of Antibodies to Conserved PfEMP1 Variants. Cell Host Microbe. 2019;26(5):579–90 e5. Epub 2019/11/15. doi: 10.1016/j.chom.2019.10.012. PubMed PMID: 31726028.

30. Lapp SA, Geraldo JA, Chien JT, Ay F, Pakala SB, Batugedara G, et al. PacBio assembly of a Plasmodium knowlesi genome sequence with Hi-C correction and manual annotation of the SICAvar gene family. Parasitology. 2018;145(1):71–84. Epub 2017/07/20. doi: 10.1017/S0031182017001329. PubMed PMID: 28720171; PubMed Central PMCID: PMCPMC5798397.

31. Lapp SA, Mok S, Zhu L, Wu H, Preiser PR, Bozdech Z, et al. Plasmodium knowlesi gene expression differs in ex vivo compared to in vitro blood-stage cultures. Malar J. 2015;14:110. Epub 2015/04/17. doi: 10.1186/s12936-015-0612-8. PubMed PMID: 25880967; PubMed Central PMCID: PMCPMC4369371.

32. Corredor V, Meyer EVS, Lapp S, Corredor-Medina C, Huber CS, Evans AG, et al. A SICAvar switching event in Plasmodium knowlesi is associated with the DNA rearrangement of conserved 3′ non-coding sequences. Molecular and Biochemical Parasitology. 2004;138(1):37–49. doi: 10.1016/j.molbiopara.2004.05.017.

33. Heather JM, Chain B. The sequence of sequencers: The history of sequencing DNA. Genomics. 2016;107(1):1–8. Epub 2015/11/12. doi: 10.1016/j.ygeno.2015.11.003. PubMed PMID: 26554401; PubMed Central PMCID: PMCPMC4727787.

34. Benavente ED, de Sessions PF, Moon RW, Grainger M, Holder AA, Blackman MJ, et al. A reference genome and methylome for the Plasmodium knowlesi A1-H.1 line. Int J Parasitol. 2018;48(3-4):191-6. Epub 2017/12/21. doi: 10.1016/j.ijpara.2017.09.008. PubMed PMID: 29258833.

35. Ahmed AM, Pinheiro MM, Divis PC, Siner A, Zainudin R, Wong IT, et al. Disease progression in Plasmodium knowlesi malaria is linked to variation in invasion gene family members. PLoS Negl Trop Dis. 2014;8(8):e3086. Epub 2014/08/15. doi: 10.1371/journal.pntd.0003086. PubMed PMID: 25121807; PubMed Central PMCID: PMCPMC4133233.

36. Ahmed MA, Quan FS. Plasmodium knowlesi clinical isolates from Malaysia show extensive diversity and strong differential selection pressure at the merozoite surface protein 7D (MSP7D). Malar J. 2019;18(1):150. Epub 2019/05/01. doi: 10.1186/s12936-019-2782-2. PubMed PMID: 31035999; PubMed Central PMCID: PMCPMC6489361.

37. Assefa S, Lim C, Preston MD, Duffy CW, Nair MB, Adroub SA, et al. Population genomic structure and adaptation in the zoonotic malaria parasite Plasmodium knowlesi. Proc Natl Acad Sci U S A. 2015;112(42):13027–32. Epub 2015/10/07. doi: 10.1073/pnas.1509534112. PubMed PMID: 26438871; PubMed Central PMCID: PMCPMC4620865.

38. Divis PCS, Duffy CW, Kadir KA, Singh B, Conway DJ. Genome-wide mosaicism in divergence between zoonotic malaria parasite subpopulations with separate sympatric transmission cycles. Mol Ecol. 2018;27(4):860–70. Epub 2018/01/03. doi: 10.1111/mec.14477. PubMed PMID: 29292549; PubMed Central PMCID: PMCPMC5918592.

39. Fong MY, Lau YL, Jelip J, Ooi CH, Cheong FW. Genetic characterisation of the erythrocyte-binding protein (PkbetaII) of Plasmodium knowlesi isolates from Malaysia. J Genet. 2019;98. Epub 2019/09/24. PubMed PMID: 31544794.

40. Gel B, Serra E. karyoploteR: an R/Bioconductor package to plot customizable genomes displaying arbitrary data. Bioinformatics. 2017;33(19):3088–90. doi: 10.1093/bioinformatics/btx346.

41. Nattestad M, Schatz MC. Assemblytics: a web analytics tool for the detection of variants from an assembly. Bioinformatics. 2016;32(19):3021–3. doi: 10.1093/bioinformatics/btw369.

42. Wang Y, Tang H, Debarry JD, Tan X, Li J, Wang X, et al. MCScanX: a toolkit for detection and evolutionary analysis of gene synteny and collinearity. Nucleic Acids Res. 2012;40(7):e49. Epub 2012/01/06. doi: 10.1093/nar/gkr1293. PubMed PMID: 22217600; PubMed Central PMCID: PMCPMC3326336.

43. Thorpe P, Escudero-Martinez CM, Cock PJA, Eves-van den Akker S, Bos JIB. Shared Transcriptional Control and Disparate Gain and Loss of Aphid Parasitism Genes. Genome Biol Evol. 2018;10(10):2716–33. Epub 2018/08/31. doi: 10.1093/gbe/evy183. PubMed PMID: 30165560; PubMed Central PMCID: PMCPMC6186164.

44. Gil M, Zanetti MS, Zoller S, Anisimova M. CodonPhyML: fast maximum likelihood phylogeny estimation under codon substitution models. Mol Biol Evol. 2013;30(6):1270–80. Epub 2013/02/26. doi: 10.1093/molbev/mst034. PubMed PMID: 23436912; PubMed Central PMCID: PMCPMC3649670.

45. Oresegun DR, Daneshvar C, Cox-Singh J. Plasmodium knowlesi – Clinical Isolate Genome Sequencing to Inform Translational Same-Species Model System for Severe Malaria. Frontiers in Cellular and Infection Microbiology. 2021;11(90). doi: 10.3389/fcimb.2021.607686.

46. Moon RW, Hall J, Rangkuti F, Ho YS, Almond N, Mitchell GH, et al. Adaptation of the genetically tractable malaria pathogen Plasmodium knowlesi to continuous culture in human erythrocytes. Proc Natl Acad Sci U S A. 2013;110(2):531–6. Epub 2012/12/26. doi: 10.1073/pnas.1216457110. PubMed PMID: 23267069; PubMed Central PMCID: PMCPMC3545754.

47. Oxford Nanopore T. Medaka: consensus sequence tool for nanopore sequences. 0.6.5 ed: Oxford Nanopore Technologies; 2019.

48. Steinbiss S, Silva-Franco F, Brunk B, Foth B, Hertz-Fowler C, Berriman M, et al. Companion: a web server for annotation and analysis of parasite genomes. Nucleic Acids Res. 2016;44(Web Server issue):W29-W34. doi: 10.1093/nar/gkw292.

49. Vaser R, Sović I, Nagarajan N, Šikić M. Fast and accurate de novo genome assembly from long uncorrected reads. Genome Res. 2017;27(5):737–46. doi: 10.1101/gr.214270.116.

50. Walker BJ, Abeel T, Shea T, Priest M, Abouelliel A, Sakthikumar S, et al. Pilon: An Integrated Tool for Comprehensive Microbial Variant Detection and Genome Assembly Improvement. PLoS ONE. 2014;9(11):e112963. doi: 10.1371/journal.pone.0112963.

51. Jain M, Koren S, Miga KH, Quick J, Rand AC, Sasani TA, et al. Nanopore sequencing and assembly of a human genome with ultra-long reads. Nat Biotechnol. 2018;36(4):338–45. Epub 2018/02/13. doi: 10.1038/nbt.4060. PubMed PMID: 29431738; PubMed Central PMCID: PMCPMC5889714.

52. Laver T, Harrison J, O’Neill PA, Moore K, Farbos A, Paszkiewicz K, et al. Assessing the performance of the Oxford Nanopore Technologies MinION. Biomol Detect Quantif. 2015;3:1–8. Epub 2016/01/12. doi: 10.1016/j.bdq.2015.02.001. PubMed PMID: 26753127; PubMed Central PMCID: PMCPMC4691839.

53. Shafin K, Pesout T, Lorig-Roach R, Haukness M, Olsen HE, Bosworth C, et al. Nanopore sequencing and the Shasta toolkit enable efficient de novo assembly of eleven human genomes. Nat Biotechnol. 2020;38(9):1044–53. Epub 2020/07/21. doi: 10.1038/s41587-020-0503-6. PubMed PMID: 32686750; PubMed Central PMCID: PMCPMC7483855.

54. Liu C, Yang X, Duffy BF, Hoisington-Lopez J, Crosby M, Porche-Sorbet R, et al. High-resolution HLA typing by long reads from the R10.3 Oxford nanopore flow cells. Hum Immunol. 2021;82(4):288–95. Epub 2021/02/23. doi: 10.1016/j.humimm.2021.02.005. PubMed PMID: 33612390.

55. Wright C. Rebasecalling of SRE and ULK GM24385 Dataset [Data Release]. EPI2ME Labs: Oxford Nanopore Technologies; 2021 [updated 18/05/2021; cited 2021 27/05/2021]. Available from: www.labs.epi2me.io/gm24385_2021.05/.

56. Divis PCS, Hu TH, Kadir KA, Mohammad DSA, Hii KC, Daneshvar C, et al. Efficient Surveillance of Plasmodium knowlesi Genetic Subpopulations, Malaysian Borneo, 2000-2018. Emerg Infect Dis. 2020;26(7):1392–8. Epub 2020/06/23. doi: 10.3201/eid2607.190924. PubMed PMID: 32568035; PubMed Central PMCID: PMCPMC7323547.

57. Diez Benavente E, Florez de Sessions P, Moon RW, Holder AA, Blackman MJ, Roper C, et al. Analysis of nuclear and organellar genomes of Plasmodium knowlesi in humans reveals ancient population structure and recent recombination among host-specific subpopulations. PLOS Genetics. 2017;13(9):e1007008. doi: 10.1371/journal.pgen.1007008.

58. Benavente ED, Gomes AR, De Silva JR, Grigg M, Walker H, Barber BE, et al. Whole genome sequencing of amplified Plasmodium knowlesi DNA from unprocessed blood reveals genetic exchange events between Malaysian Peninsular and Borneo subpopulations. Sci Rep. 2019;9(1):9873. Epub 2019/07/10. doi: 10.1038/s41598-019-46398-z. PubMed PMID: 31285495; PubMed Central PMCID: PMCPMC6614422.

59. International Human Genome Sequencing C, Lander ES, Linton LM, Birren B, Nusbaum C, Zody MC, et al. Initial sequencing and analysis of the human genome. Nature. 2001;409(6822):860–921. doi: 10.1038/35057062.

60. Li H. Minimap2: pairwise alignment for nucleotide sequences. Bioinformatics. 2018;34(18):3094–100. doi: 10.1093/bioinformatics/bty191.

61. Li H. A statistical framework for SNP calling, mutation discovery, association mapping and population genetical parameter estimation from sequencing data. Bioinformatics (Oxford, England). 2011;27(21):2987–93. doi: 10.1093/bioinformatics/btr509.

62. Li H, Handsaker B, Wysoker A, Fennell T, Ruan J, Homer N, et al. The Sequence Alignment/Map format and SAMtools. Bioinformatics. 2009;25(16):2078–9. doi: 10.1093/bioinformatics/btp352.

63. Quinlan AR, Hall IM. BEDTools: a flexible suite of utilities for comparing genomic features. Bioinformatics. 2010;26(6):841–2. doi: 10.1093/bioinformatics/btq033.

64. Kolmogorov M. Fast and accurate de novo assembler for single molecule sequencing reads: fenderglass/Flye. 2019.

65. Laetsch DR, Blaxter ML. BlobTools: Interrogation of genome assemblies. F1000Research. 2017;6:1287. doi: 10.12688/f1000research.12232.1.

66. Morgulis A, Coulouris G, Raytselis Y, Madden TL, Agarwala R, Schäffer AA. Database indexing for production MegaBLAST searches. Bioinformatics. 2008;24(16):1757–64. doi: 10.1093/bioinformatics/btn322.

67. Hunt M, Silva ND, Otto TD, Parkhill J, Keane JA, Harris SR. Circlator: automated circularization of genome assemblies using long sequencing reads. Genome Biol. 2015;16:294. doi: 10.1186/s13059-015-0849-0.

68. Flynn JM, Hubley R, Goubert C, Rosen J, Clark AG, Feschotte C, et al. RepeatModeler2 for automated genomic discovery of transposable element families. Proc Natl Acad Sci U S A. 2020;117(17):9451–7. Epub 2020/04/18. doi: 10.1073/pnas.1921046117. PubMed PMID: 32300014; PubMed Central PMCID: PMCPMC7196820.

69. Kohany O, Gentles AJ, Hankus L, Jurka J. Annotation, submission and screening of repetitive elements in Repbase: RepbaseSubmitter and Censor. BMC Bioinformatics. 2006;7(1):474. doi: 10.1186/1471-2105-7-474.

70. Fu L, Niu B, Zhu Z, Wu S, Li W. CD-HIT: accelerated for clustering the next-generation sequencing data. Bioinformatics (Oxford, England). 2012;28(23):3150–2. doi: 10.1093/bioinformatics/bts565.

71. Li W, Godzik A. Cd-hit: a fast program for clustering and comparing large sets of protein or nucleotide sequences. Bioinformatics (Oxford, England). 2006;22(13):1658–9. doi: 10.1093/bioinformatics/btl158.

72. Bailly-Bechet M, Haudry A, Lerat E. “One code to find them all”: a perl tool to conveniently parse RepeatMasker output files. Mobile DNA. 2014;5(1):13. doi: 10.1186/1759-8753-5-13.

73. Ellinghaus D, Kurtz S, Willhoeft U. LTRharvest, an efficient and flexible software for de novo detection of LTR retrotransposons. BMC Bioinformatics. 2008;9:18. doi: 10.1186/1471-2105-9-18.

74. Gremme G, Steinbiss S, Kurtz S. GenomeTools: a comprehensive software library for efficient processing of structured genome annotations. IEEE/ACM Trans Comput Biol Bioinform. 2013;10(3):645–56. Epub 2013/10/05. doi: 10.1109/TCBB.2013.68. PubMed PMID: 24091398.

75. Alonge M, Soyk S, Ramakrishnan S, Wang X, Goodwin S, Sedlazeck FJ, et al. RaGOO: fast and accurate reference-guided scaffolding of draft genomes. Genome Biol. 2019;20(1):224. Epub 2019/10/30. doi: 10.1186/s13059-019-1829-6. PubMed PMID: 31661016; PubMed Central PMCID: PMCPMC6816165.

76. Stanke M, Keller O, Gunduz I, Hayes A, Waack S, Morgenstern B. AUGUSTUS: ab initio prediction of alternative transcripts. Nucleic Acids Res. 2006;34(Web Server issue):W435-W9. doi: 10.1093/nar/gkl200.

77. Simão FA, Waterhouse RM, Ioannidis P, Kriventseva EV, Zdobnov EM. BUSCO: assessing genome assembly and annotation completeness with single-copy orthologs. Bioinformatics. 2015;31(19):3210–2. doi: 10.1093/bioinformatics/btv351.

78. Okonechnikov K, Conesa A, García-Alcalde F. Qualimap 2: advanced multi-sample quality control for high-throughput sequencing data. Bioinformatics. 2016;32(2):292–4. doi: 10.1093/bioinformatics/btv566.

79. Cabanettes F, Klopp C. D-GENIES: dot plot large genomes in an interactive, efficient and simple way. PeerJ. 2018;6:e4958. Epub 2018/06/12. doi: 10.7717/peerj.4958. PubMed PMID: 29888139; PubMed Central PMCID: PMCPMC5991294.

80. Ren J, Chaisson MJP. lra: the Long Read Aligner for Sequences and Contigs. preprint. Bioinformatics, 2020 2020/11/17/. Report No.

81. Pedersen BS, Quinlan AR. Mosdepth: quick coverage calculation for genomes and exomes. Bioinformatics. 2018;34(5):867–8. doi: 10.1093/bioinformatics/btx699.

82. Jiang T, Liu Y, Jiang Y, Li J, Gao Y, Cui Z, et al. Long-read-based human genomic structural variation detection with cuteSV. Genome Biol. 2020;21(1):189. doi: 10.1186/s13059-020-02107-y.

83. Jeffares DC, Jolly C, Hoti M, Speed D, Shaw L, Rallis C, et al. Transient structural variations have strong effects on quantitative traits and reproductive isolation in fission yeast. Nature Communications. 2017;8(1):14061. doi: 10.1038/ncomms14061.

84. Thorvaldsdóttir H, Robinson JT, Mesirov JP. Integrative Genomics Viewer (IGV): high-performance genomics data visualization and exploration. Briefings in Bioinformatics. 2013;14(2):178–92. doi: 10.1093/bib/bbs017.

85. Emms DM, Kelly S. OrthoFinder: phylogenetic orthology inference for comparative genomics. Genome Biol. 2019;20(1):238. Epub 2019/11/16. doi: 10.1186/s13059-019-1832-y. PubMed PMID: 31727128; PubMed Central PMCID: PMCPMC6857279.

86. Edgar RC. MUSCLE: multiple sequence alignment with high accuracy and high throughput. Nucleic Acids Res. 2004;32(5):1792–7. Epub 2004/03/23. doi: 10.1093/nar/gkh340. PubMed PMID: 15034147; PubMed Central PMCID: PMCPMC390337.

87. Cock PJ, Antao T, Chang JT, Chapman BA, Cox CJ, Dalke A, et al. Biopython: freely available Python tools for computational molecular biology and bioinformatics. Bioinformatics. 2009;25(11):1422–3. Epub 2009/03/24. doi: 10.1093/bioinformatics/btp163. PubMed PMID: 19304878; PubMed Central PMCID: PMCPMC2682512.

88. Capella-Gutierrez S, Silla-Martinez JM, Gabaldon T. trimAl: a tool for automated alignment trimming in large-scale phylogenetic analyses. Bioinformatics. 2009;25(15):1972–3. Epub 2009/06/10. doi: 10.1093/bioinformatics/btp348. PubMed PMID: 19505945; PubMed Central PMCID: PMCPMC2712344.

89. Eves-van den Akker S, Laetsch DR, Thorpe P, Lilley CJ, Danchin EG, Da Rocha M, et al. The genome of the yellow potato cyst nematode, Globodera rostochiensis, reveals insights into the basis of parasitism and virulence. Genome Biol. 2016;17(1):124. Epub 2016/06/12. doi: 10.1186/s13059-016-0985-1. PubMed PMID: 27286965; PubMed Central PMCID: PMCPMC4901422.

90. Thorpe P, Escudero-Martinez CM, Eves-van den Akker S, Bos JIB. Transcriptional changes in the aphid species Myzus cerasi under different host and environmental conditions. Insect Mol Biol. 2020;29(3):271–82. Epub 2019/12/18. doi: 10.1111/imb.12631. PubMed PMID: 31846128; PubMed Central PMCID: PMCPMC7317760.

